# Synthetic maize centromeres transmit chromosomes across generations

**DOI:** 10.1101/2022.09.19.508542

**Authors:** R. Kelly Dawe, Jonathan I. Gent, Yibing Zeng, Han Zhang, Fang-Fang Fu, Kyle W. Swentowsky, Dong won Kim, Na Wang, Jianing Liu, Rebecca D. Piri

**Affiliations:** Department of Genetics, University of Georgia, Athens GA 30602, USA; Department of Plant Biology, University of Georgia, Athens GA 30602, USA; Institute of Bioinformatics, University of Georgia, Athens, GA 30602; Co-Innovation Center for Sustainable Forestry in Southern China, Nanjing Forestry University, Nanjing, China 210037

## Abstract

Centromeres are long, often repetitive regions of genomes that bind kinetochore proteins and ensure normal chromosome segregation. Engineering centromeres that function in vivo has proven to be difficult. Here we describe a LexA-CENH3 tethering approach that activates functional centromeres at maize synthetic repeat arrays containing LexO binding sites. The synthetic centromeres are sufficient to cause chromosome breakage and release of chromosome fragments that are passed through meiosis and into progeny. Several independent chromosomes were identified, each with newly created centromeres localized over the repeat arrays where they were directed. The new centromeres were self-sustaining and stably transmitted chromosomes to progeny in the absence of the LexA-CENH3 activator. Our results demonstrate the feasibility of using synthetic centromeres for karyotype engineering applications.

## Introduction

Decades of advances in genetics have led to a strong awareness that traits related to the most pressing problems in agriculture are polygenic. Information about the combined effects of hundreds of genes known to be involved in drought, heat, and pathogen pressure (*1*) might be used to substantially improve plant performance, but we lack the methods necessary to genetically engineer crops at this scale. Many have suggested that synthetic biology approaches could be used to meet this challenge (e.g. (*2*–*4*)). In bacteria and yeast, it is possible to chemically synthesize megabase-scale sequences and shuttle them between species to alter gene order and structure (*4*–*7*). Similar methods have been used to move DNA from yeast into human cells (*4*) and could be adapted to plants as well (*8*). Engineering small chromosomes for practical applications in medicine and agriculture seems a reasonable near-term goal (*8*–*10*).

A major obstacle in whole chromosome design is the synthesis of functional centromeres – sites of kinetochore assembly that are required for chromosome segregation. Centromeres can span hundreds of kilobases of repetitive, ill-defined sequence. Centromere specification is also strongly epigenetic, such that the DNA sequence matters less than the centromere proteins that are stably bound to the sequence (*11*). Early studies established that long tracts of native centromeric DNA can form artificial chromosomes in human cell lines (*12*), but success rates are low and rely on the presence of a primate-specific centromere protein (CENP-B) (*13*). Many have suggested that centromere engineering might be better accomplished via protein tethering (*13*–*19*). In this approach, a cell line is first transformed with an array of LacO DNA binding sites that provide a platform to build a new centromere. The line is then transformed with a gene that encodes a fusion between the corresponding DNA binding protein (LacI) and one of several proteins that associate with native centromeres (*14, 15, 17, 20*). Bound by the fusion proteins, the synthetic DNA recruits additional kinetochore proteins and shows features of a functional centromere (*14, 15, 17, 20*). The strengths of the tethering approach are that it relies on defined components and is broadly applicable to all species.

The most effective synthetic centromeres have been based on recruiting CENP-A/CENH3 to LacO arrays. CENP-A/CENH3, a histone H3 variant, is the defining feature of centromere-specific nucleosomes (*21*). In humans and Drosophila, CENP-A is deposited by specialized chaperones called HJURP and Cal1 (*22, 23*). When LacI-CENP-A, LacI-HJURP, or LacI-Cal1 fusion proteins were introduced into cell lines carrying LacO arrays on chromosome arms, active centromeres formed over the LacO repeats (*16, 20, 23*). The LacO sites (and often flanking sequences) retained kinetochore activity during mitosis even after the LacI fusion proteins were removed (*18*–*20*). These data established that it is only necessary to seed centromere proteins at an ectopic site to activate the self-replication mechanism. The first effort to test synthetic centromeres in living organisms was recently carried out using Drosophila (*19*). The authors inserted LacO arrays into several locations of the Drosophila genome and observed centromere activation by Cal1-CENP-A tethering at a high frequency in developing animals. However, the resulting chromosome instability led to apoptosis (a cell death response unique to animals) and loss of lineages containing the new centromeres (*19*), making it difficult to interpret whether the newly activated centromeres could independently control chromosome segregation or be transmitted through meiosis into progeny.

### Molecular characterization of the synthetic centromere platform ABS4

We previously prepared long synthetic repeat arrays composed of 157 bp monomers that contain binding sites for several known DNA binding proteins (*24*). Each monomer contains binding motifs for *Escherichia coli* LacI and LexA and budding yeast Gal4. The arrays were biolistically transformed into maize and recovered in three locations on different chromosome arms. The longest of these is Arrayed Binding Sites on chromosome 4 (ABS4) which is composed of a mixture of 157-bp ABS monomers, a marker plasmid called pAHC25, and possibly other rearranged genomic sequences distributed over a ∼1,100 kb region (*24*). For this study, we sequenced a line containing ABS4 to determine the location of the array and copy number of ABS monomers within it. The data suggest that ABS4 contains ∼2,400 copies of ABS (about 377 kb) and ∼10.5 copies of pAHC25 (about 101 kb)(Figure 1A). We detected two junctions between the transformed sequences and chromosome 4, one at position 183583798 (with ABS) and another at position 185020754 (with pAHC25). This location corresponds to the long arm of chromosome 4, roughly 44 cM from the native centromere.

**Figure 1.**
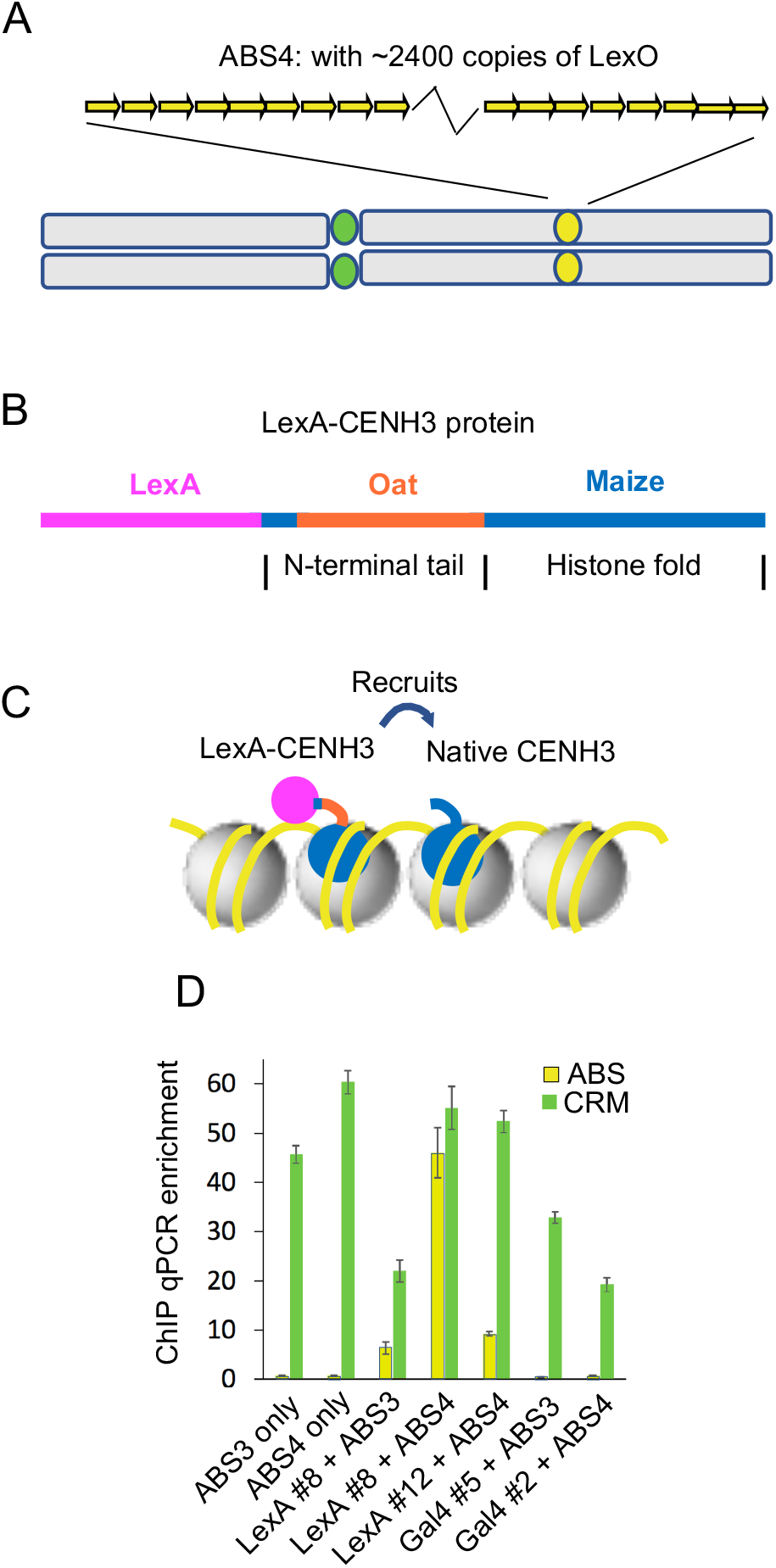
LexA-CENH3(v2) recruits native CENH3 to ABS4. **A**) Relative positions of centromere 4 (yellow) and ABS4 (green) on chromosome 4. **B**) Schematic of the the LexA-CENH3 chimeric protein. **C)** Cartoon illustrating how LexA-CENH3 may interact with DNA and nucleosomes in ABS arrays. Not shown are other constitutive centromere proteins that bind to CENH3, some of which recruit additional CENH3 to maintain stable centromere positions. **D**) CENH3 ChIP enrichment relative to input, measured by qPCR in plants with v1 constructs. LexA-CENH3(v1) is abbreviated to LexA and Gal4-CENH3(v1) is abbreviated to Gal4. Each ChIP was from an individual plant that was heterozygous ABS only or ABS plus a CENH3 fusion transgene. Two different transgenic events for each construct were tested (noted with #) along with ABS3 and ABS4. The CRM retroelement is common in native centromeres and is used as a positive control. Y-axis indicates qPCR abundance using ABS- and CRM2-specific primers. Error bars are standard deviations from qPCR technical replicates.

### ChIP assays for centromere activity at ABS using different CENH3 fusions

In Drosophila and human, centromere protein tethering has made use of the LacI DNA binding domain (*14*–*20*). We opted against using LacI because of its large size (366 aa), as data from Arabidopsis suggested that large N-terminal additions to CENH3 could interfere with function (*25*). We prepared constructs to test the efficacy of the Gal4 and the LexA DNA binding domains when fused to the N-terminus of maize CENH3. In each case, we also replaced the native N-terminal tail of maize CENH3 with the N-terminal tail from oat CENH3 (Figure 1B). This modification removed the epitope for an maize CENH3 antibody (*26*) and added the epitope for an oat CENH3 antibody (*27*). Removing the maize epitope from the transgene allowed us to test whether the transgene could recruit native CENH3 to the ABS4 array (Figure 1C). Our first generation (v1) constructs were composed of a ∼1.7 kb segment of the native *CENH3* promoter driving the chimeric proteins as intronless open reading frames. The constructs were transformed into maize, crossed to lines carrying ABS loci (either ABS4 or ABS3 on chromosome 3 (*24*)), and the heterozygous progeny subjected to Chromatin ImmunoPrecipitation (ChIP) using the maize specific CENH3 antibody. The results were assayed by both ChIP-qPCR and ChIP-seq (Figure 1D, Figure S1). *LexA-CENH3(v1*) but not *Gal4-CENH3(v1*) showed significant enrichment of ABS (Figure 1D). These results demonstrate that *LexA-CENH3(v1*) interacts with ABS and recruits native CENH3 to ABS sequences.

### Reengineering of the LexA-CENH3 transgene and demonstration of whole plant functionality

Despite the strong ChIP enrichment of ABS in leaf tissue, the *LexA-CENH3(v1*) transgene was only weakly expressed in meiotic cells and did not rescue the embryo lethal phenotype of a homozygous *cenh3* null mutant (Figure S2). We had previously established that a ∼5 kb region of the *CENH3* genomic sequence with a ∼2.1 kb promoter and all introns could complement the *cenh3* null allele (*28*). We modified this construct by inserting LexA and oat sequences into the *CENH3* coding sequence while retaining the order of and sequence of introns (*LexA-CENH3(v2)*). Plants carrying the *LexA-CENH3(v2*) transgene and homozygous for the *cenh3* null allele were viable (Figure S2), although three of the four plants were thin and weak. The fourth plant was vigorous; it was self-crossed and proved to be homozygous for *LexA-CENH3(v2*) (and *cenh3*/*cenh3*). Protein immunoblot analysis of this plant demonstrated that native maize CENH3 protein was absent, while the larger LexA-CENH3(v2) protein, identified by the oat CENH3 epitope, was readily detectable (Figure S3). These results show that LexA-CENH3(v2) can substitute for native CENH3, is properly expressed in all tissues, and does not interfere with the assembly of functional kinetochores.

### Chromosome breakage in somatic tissues of ABS4/LexA-CENH3 lines

Barbara Mclintock was the first to describe the cytological and genetic behavior of dicentric chromosomes (*29, 30*). When located far apart on a chromosome, two centromeres are just as likely to move to opposite poles on the spindle as they are to move to the same pole (Figure 2A). The opposite orientation results in criss-cross patterns with chromatin bridges that can rupture in the spindle midzone, creating double stranded DNA breaks (*31, 32*). The two breaks can then fuse (by non-homologous end joining) in the next cell cycle to recover the original dicentric condition, leading to repeated cycles of breakage and rejoining (*30*). Another outcome is that the breaks can heal by acquiring telomeres, separating the previously dicentric chromosome into two independent chromosomes (*30, 33*). In a minority of cases, the bridge is pulled to one pole without breaking (*31, 34*).

**Figure 2.**
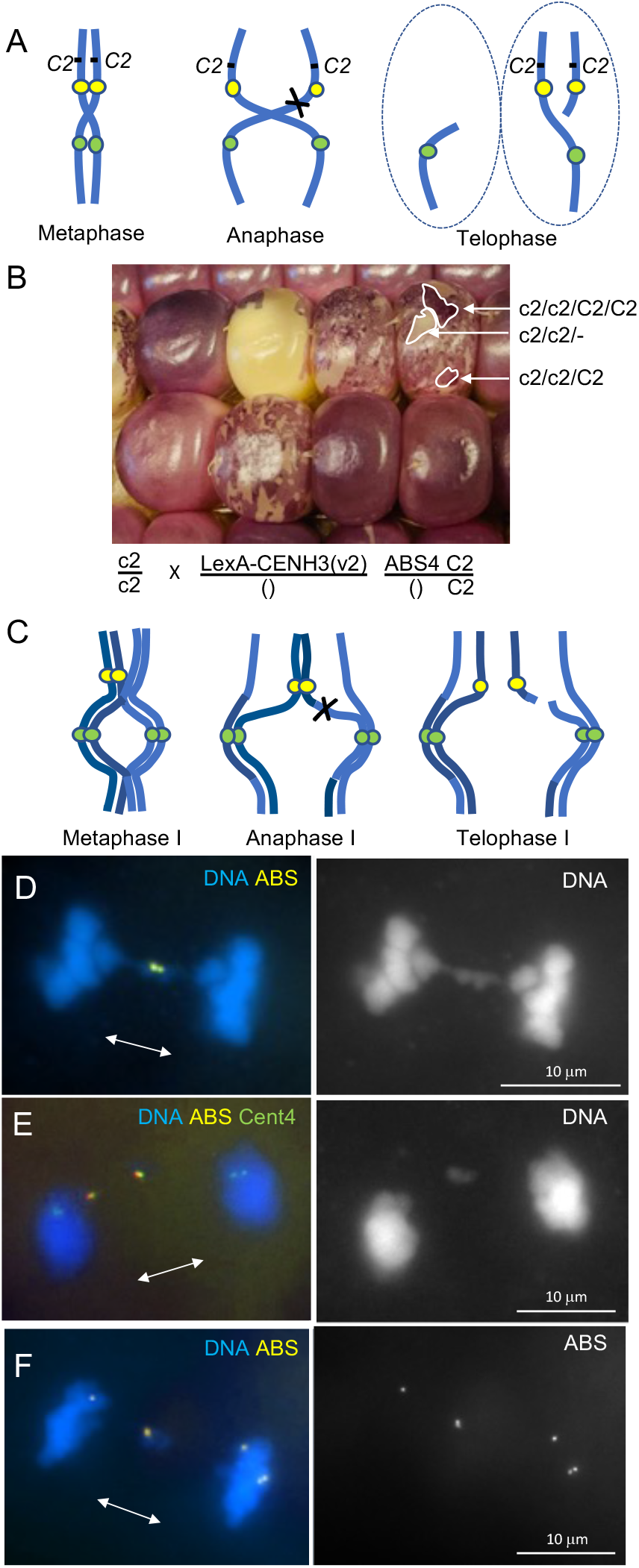
Segregation errors in LexA-CENH3 ABS4 plants. **A**) Illustration of dicentric chromosome segregation where only one chromosome breaks, creating a daughter cell with no ABS4 (or the closely linked *C2* locus) and a daughter cell with two copies of ABS4 (and two copies of *C2*). ABS4 is shown in yellow and Cent4 in green. Homologous sister chromatids without ABS4 (not shown) are expected to segregate properly. **B**) Kernels from a cross where two copies of chromosome 4 (with *c2* null alleles) came from the female, and LexA-CENH3 and one copy of chromosome 4 with ABS4 and *C2* came from the male (the cross is shown below the image). Kernels that received ABS4 and LexA-CENH3 are sectored, showing patches of colorless and deeply pigmented tissue. The patterns are consistent with the model in A and cytological observations of roots grown from sectored kernels (Figure S4). **C**) A model of how sister centromere cohesion can restrain ABS4 to form a bridge and cause chromosome breakage if there is recombination between the native centromere and ABS. The parental chromosomes are in slightly different shades of blue to illustrate recombination. **D**) Anaphase I showing a bridge with ABS at the connecting point. Right image shows DNA. **E**) Telophase I cell showing breakage and release of an ABS-containing fragment in the midzone. Cent4 is labeled green (one on each sister centromere) showing that ABS is separated from the native centromere. Right image shows DNA. **F**) Late anaphase I cell showing six ABS loci, where two are expected. The accumulation of ABS loci must have occurred prior to meiosis. Right image shows ABS loci.

The ABS4 locus is about 12 cM from the *colorless2* (*C2*) gene that is required for purple pigmentation in the outer cell layers (aleurone) of the starchy part of the seed (endosperm). Unlike the embryo, the endosperm is triploid, where two complete genomes are contributed by the female and one from male. The intensity of the purple color conferred by a wild-type *C2* allele is dosage dependent, such that the seeds are colorless in the absence of *C2*, one copy of *C2* gives a faint color and two copies a darker color (*35*). In control crosses between a line homozygous for a recessive mutant *c2* allele (*c2/c2*) as a female and a wild-type (*C2/C2*) line as a male, the seeds are a uniformly faint purple color. Likewise, the seeds were uniformly pigmented when we crossed *c2/c2* lines by *C2/C2* plants carrying ABS4 or *LexA-CENH3(v2*) alone.

When we crossed *c2/c2* females by *C2/C2* males heterozygous for both *LexA-CENH3(v2*) and ABS4, we observed sectors of colorless, pale, and deeply pigmented tissue juxtaposed on endosperm (Figure 2B). Since the *C2* locus is on the distal side of ABS4 (towards the telomere), the color alterations are best explained by segregation defects associated with activation of ABS4 to become a second functional centromere. One explanation for the frequent sectors is that the dicentric chromosome often segregates to one pole without breaking, perhaps because the newly formed ABS4 centromeres are relatively weak compared to the native centromere. If one chromatid breaks and the other does not, one of the two daughter cells will lack *C2* (and be colorless) and the other will contain two copies of *C2* (and be a dark color). In support of this view, we grew roots from sectored seeds and observed that about 4.0% of the nuclei lack ABS and 5.5% have two copies of ABS (Figure S4A).

We also observed that plants heterozygous for *LexA-CENH3(v2*) and ABS4 tended to be short and have asymmetric leaves (Figure S5), suggesting that the loss and gain of genes on chromosome 4L negatively impacted plant development. Similar phenotypes were observed in pea plants carrying a dicentric chromosome (*33*).

### Chromosome breakage in meiotic cells of ABS4/LexA-CENH3 lines

In meiosis I, criss-cross patterns are not expected because the sister centromeres remain attached throughout the first division (*36*). Nevertheless, because ABS4 is ∼44 cM from the native centromere, recombination will frequently place one copy of ABS4 on each chromosome (Figure 2C).

When this occurs, sister centromere cohesion will prevent complete disjunction, resulting in bridge structures where the arms remain attached specifically at the ABS locus. Bridges of this type were readily observed at mid-anaphase I (Figure 2D). An analysis of 68 mid-anaphase I cells revealed that 63% of the recombinant chromosomes remained connected at ABS4 (Figure S4B). At late anaphase I, the bridges were resolved by pulling forces, either by breaking the cohesin bonds or by breaking one of the chromosome arms. Multiple examples of broken, ABS-containing fragments were observed in the midzone of late anaphase-telophase cells. A probe specific to centromere 4 (Cent4 (*37*)) was used to demonstrate that ABS had broken away from its native centromere in many cases (Figure 2E). We also observed meiosis I cells with no ABS or multiple ABS loci caused by mitotic events that occurred during the preceding mitoses (Figure 2F). However, few if any new fragments were produced during meiosis II, which appeared to proceed normally with no evidence of anaphase II bridges. These results establish that LexA-CENH3-activated centromeres are of sufficient strength to break off fragments of chromosome 4L that are visible in meiosis.

### Identification of ABS-driven neochromosomes

Although many of the fragments we observed in meiosis were newly broken or showing aberrant segregation and lost, many more were likely included in telophase nuclei where they could be transmitted to the next generation. The fragments presumably also carried CENH3 or other chromatin marks that could enable ABS4 to serve as the centromere for a new chromosome (neochromosome). However, any neochromosome derived from chromosome 4L will lack hundreds of genes that are normally contributed by chromosome 4S. Since many of those genes are required for gametophyte development (*38*), such fragments cannot be transmitted to the next generation unless they are accompanied by a normal chromosome 4, which rescues the deficiency but also adds a second copy of 4L. The changes in gene dosage caused by adding an extra chromosome arm can likewise be detrimental to gametophytes, particularly through the male because pollen tubes compete with each other to reach the egg cell.

Assuming the neochromosome can be successfully passed through gametophytes, the resulting progeny will be partially trisomic. Trisomic maize plants usually survive to maturity, but three chromosomes cannot pair properly in meiosis, causing errors in segregation that can further reduce the likelihood of passing on the chromosome (*39*). These factors can make it difficult to recover and propagate neochromosomes. Nevertheless, we have identified independent ABS chromosomes using two different approaches.

The first was discovered in the progeny of a cross between a *c2*/*c2* female and a *C2*/*C2* male heterozygous for both *LexA-CENH3(v2*) and ABS4. We noticed that some kernels were sectored but lacked *LexA-CENH3(v2*), suggesting that ABS had acquired independent centromere activity. One such kernel was planted and crossed to *c2*/*c2* in both directions, where it transmitted purple kernels (carrying *C2*) at a low frequency of 7% through the female and 11% through the male. However, most of the seedlings that grew from purple kernels did not have ABS when scored by PCR (14/17), suggesting that the ABS-containing chromosome was unstable. One plant that retained ABS was crossed again in both directions, where it showed improved transmission of C2 (35% and 22% respectively). Root tips from the progeny of these crosses were analyzed cytologically and found to be partially trisomic, with two copies of normal chromosome 4 and a fragment chromosome with ABS that we refer to as Neochromosome 4L number 1 (Neo4L-1) (Figure 3A).

**Figure 3.**
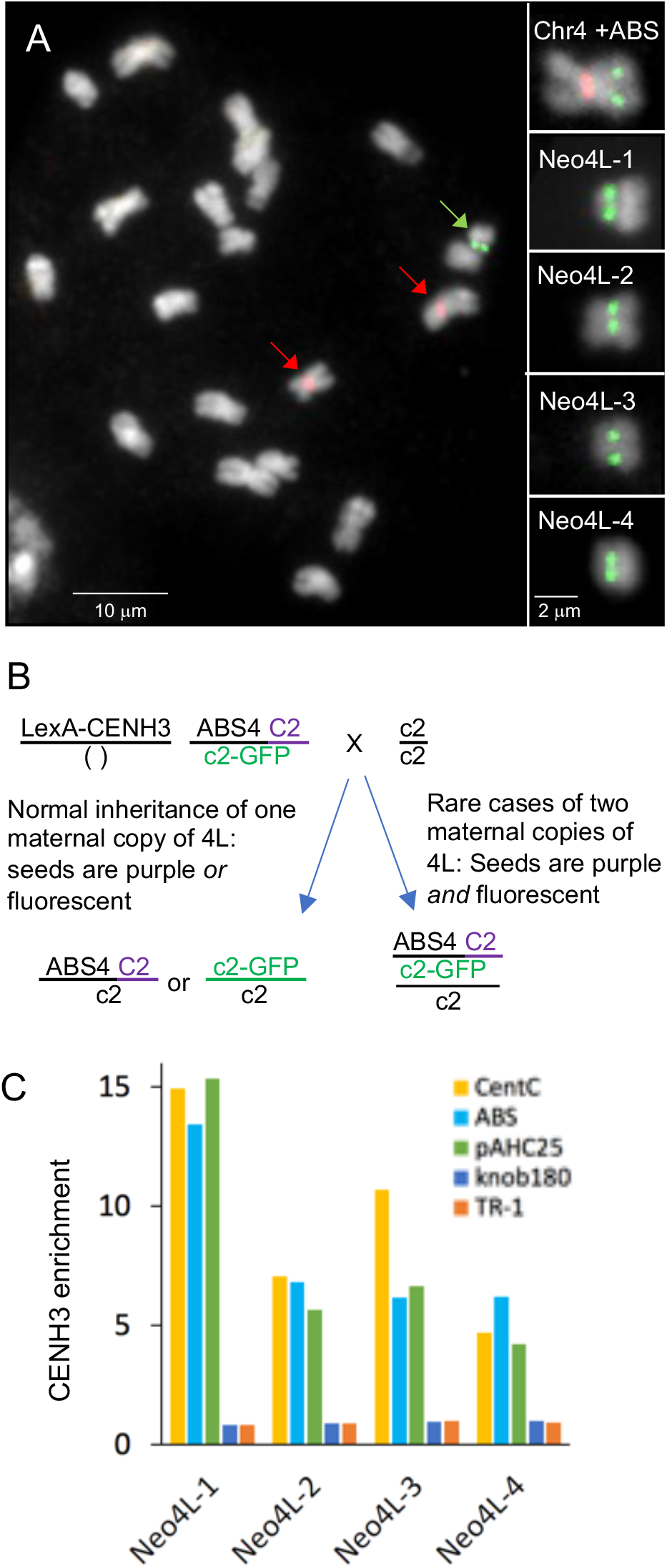
Analysis of Independent chromosomes 4L. **A**) Chromosome 4L neochromosomes. The main panel shows a full karyotype and typical partial trisomy of these lines, with two copies of normal chromosome 4 (red arrows) and Neo4L-1 (green arrow). To the right are images of the intact ABS4-containing chromosome and four neochromosomes. Neo4L-1 has a knob on the arm that makes it appear larger than the others. Cent4 is labeled in red and ABS in green. **B**) The cross used to identify Neo4L-1, Ne4L-2 and Neo4L-3. Purple/fluorescent kernels should be partially trisomic. **C**) CENH3 CUT&Tag results from each neochromosome showing enrichment for ABS and pAHC25. CentC is a native centromere repeat, knob180 and TR-1 are non-centromeric repeats.

Three other neochromosomes were identified in a screen for seeds that had received two copies of chromosome 4L from the *LexA-CENH3(v2*) ABS4 parent. For this experiment, we obtained a fluorescent null allele of *c2* (*c2-GFP*) from a public reverse genetics resource (*40*). The *c2-GFP* allele has a stable *Dissociation* transposable element containing GFP inserted in the second exon of the *c2* allele. Plants of the genotype ABS4 *C2*/*c2-GFP LexA-CENH3(v2*) were crossed as females to a *c2/c2* line (Figure 3B) and screened for kernels that were both purple and fluorescent (Figure S6). We observed 181 kernels that were both purple and fluorescent in a sample of 16,757 kernels, indicating that as many as 1% of the kernels from ABS lines may have potential neochromosomes (Table S1). To test if the screen was working as intended, we first planted a sample of 3 purple/fluorescent seeds and then another sample of 20 purple/fluorescent seeds (Table S1). Of these 23 plants, 17 lacked ABS, suggesting that if neochromosomes were present they were unstable. Two of the six plants that retained ABS grew poorly and could not be crossed, and four others were crossed in both directions to *c2/c2*. One showed nearly Mendelian segregation of *C2* and did not have a visible neochromosome (this may be a translocation). Three others transmitted *C2* at low frequencies, and progeny contained partially trisomic individuals with two copies of normal chromosome 4 and ABS-containing neochromosomes that we named Neo4L-2, Neo4L-3 and Neo4L-4 (Figure 3A).

Each of the four chromosomes displayed cytological constrictions at the ABS4 array (Figure 3A), which is a classic indicator of centromere location. To confirm that centromeres had formed over ABS4, we carried out CUT&Tag (*41*), a method that can be used to localize CENH3 on chromatin extracted from small amounts of leaf tissue. The data show that CENH3 is localized over ABS and pAHC25 on all four chromosomes (Figure 3C), although the enrichment for ABS and native centromere sequence CentC was relatively low in this differentiated tissue. We went on to carry out CUT&Tag on an immature ear from a Neo4L-1 plant (with a high proportion of dividing cells). The ear data showed a ∼97-fold enrichment of ABS sequences, exceeding the enrichment observed for the native centromere repeat CentC (Figure S7). CENH3 was also localized over a ∼350 kb portion of chromosome 4L adjacent to the junction with pAHC25 at position 185020305 on the reference genome, demonstrating that at least for Neo4L-1, CENH3 has spread into flanking chromatin.

### Genetic properties of neochromosomes

Transmission data are available for all four neochromosomes (Table S2). Neo4L-1, Neo4L-3 and Neo4L-4 were discovered in plants that lacked *LexA-CENH3(v2*), indicating that the neochromosomes had acquired stable centromeres very quickly after meiosis. The first generation Neo4L-2 plant carried *LexA-CENH3(v2*), but we chose plants that lacked *LexA-CENH3(v2*) for crossing in the second generation. The data are similar for all neochromosomes, showing 21-31% transmission through the female and variable frequencies through the male (0-22%, likely being influenced by how much pollen was applied, since excess pollen increases the amount of pollen competition and could select against partial trisomies) (Table S2). These results demonstrate that after a synthetic centromere is activated and stabilized during plant growth, the LexA-CENH3 transgene is not necessary for future transmission.

Although all four neochromosomes were successfully passed through meiosis, none were completely stable in somatic tissues during the period of this study. When the neochromosomes (marked by *C2*) were passed through the male to a *c2*/*c2* line, there were invariably kernels with colorless sectors, which we interpret as occasional chromosome loss. We also observed that about 31% of the seedlings grown from *C2* seeds lacked ABS4 when genotyped (Table S2). This result is not a result of genetic recombination between ABS4 and *C2*. We assayed 87 colorless seeds from trisomic plants segregating neochromosomes and found only one with ABS, or about ∼1% recombination (Table S2). Instead, the loss of neochromosomes in embryos appears similar to a phenomenon known as centromere-mediated chromosome elimination that is observed in crosses between lines with widely divergent centromere sizes (or CENH3 structure) (*25, 28, 42*). Centromere-mediated chromosome elimination occurs primarily in embryos (and to a lesser extent in endosperm (*43*)) and rarely exceeds a frequency of ∼45% (*25*), which mirrors what we have observed with neochromosome loss in early generations.

## Discussion

Discussions of whole genome synthetic biology often leave out the issue of recreating centromeres, which are hundreds of kilobases long and generally do not function properly unless they are bound to CENP-A/CENH3 and/or other constitutive centromere proteins. Less appreciated is the flip side of this observation, which is that apparently any non-genic sequence has the potential to become centromeric (*44, 45*). Even a short array of bacterial LacO repeats can cause centromere movement when tethered to one of several constitutive centromere proteins (*14, 15, 17, 20*). Using a LexA-based system in maize, we have demonstrated that the basic principles behind targeting CENP-A/CENH3 to a synthetic repeat array to activate mitotically stable centromeres (*18*–*20*) are applicable to the plant kingdom. We have further demonstrated that centromeres engineered in this way can become permanent features of a stably modified genome. Newly created maize synthetic centromeres can drive independent chromosome segregation over several generations in the absence of the original activator protein.

On native centromeres, CENH3 is sparsely distributed over regions as long as 2 Mb (*46*), but there are examples of centromeres that have shifted to new locations that are ∼300-500 kb (*45, 47*)). Here we recruited CENH3 to a region that may be as large as 800 kb (∼377 kb of ABS, 101 kb of pAHC25, and 350 kb of genomic DNA). Yet, as we show here, providing a large binding platform does not assure stable chromosome inheritance, particularly immediately after a centromere is formed. The limited data from our genetic screen suggests that only ∼13% (3/23) of the ABS-containing fragments that are passed through meiosis will become neochromosomes in progeny. We also observed neochromosome loss in later generations, which may reflect the activity of a centromere size surveillance mechanism (*28, 42*), or other structural abnormalities associated with centromeres or telomeres in these newly formed chromosomes.

With the means to engineer synthetic centromeres, it should now be possible to determine with more clarity the limits of centromere size in maize and other species. What is known so far indicates that functional centromere size is positively correlated with genome size (*48, 49*). Our success with a large genome plant such as maize bodes well for success in smaller genome plants such as *Setaria viridis* or *Arabidopsis thaliana*, or algal species such as *Chlamydomonas reinhardtii*. It may also be possible to extend our general approach to create circular plant artificial chromosomes similar to the artificial chromosomes built by CENP-A tethering in animal cell lines (*18*). When paired with continued efforts to improve large molecule transformation in plants, synthetic centromeres could become important tools for addressing global challenges related to the bioeconomy and food security.

## Methods

### Molecular characterization of ABS4

To create the ABS4 locus, amplified arrays of ABS monomers and the selectable marker plasmid pAHC25 (*50*) were transformed into maize using biolistic transformation (*24*). The location of ABS4 was previously known only by cytological position. To determine the exact location of the ABS4 array, we performed paired-end 150 Illumina sequencing at 16X coverage of an ABS4 heterozygous line. pAHC25 and ABS monomer sequences were concatenated with the B73 genome to create a reference for mapping. The reads were aligned with the Burrows-Wheeler Aligner (*51*) and the discordant and soft-clipped alignments in pAHC25 and ABS sequences were screened (by eye) using the Integrative Genomics Viewer (IGV). The coordinates of reads mapped to B73 were manually identified in the discordant read pair and clustered them based on their position. We then carried *de novo* assembly with discordant read pairs from individual clusters using SPAdes (v3.10.0). Upon the generation of consensus sequences, we identified the precise junction points by Blasting (v2.2.26) the assembled short segments against the reference sequence. By this means, we identified two junctions on chromosome 4: one between pAHC25 and Chr4 position at position 185020754 and another between ABS and Chr4 position 183583798 (in Zm-B73-REFERENCE-NAM-5.0 coordinates). Primers were developed to detect the second polymorphism for genotyping. One pair, abs4-p1 (TACCCTGGTTAGAGGGAGCC) and abs4-p4 (AGCCAGGCGGATAGAAGC) amplify the wild-type locus. Two pairs amplify the ABS insertion: abs4-p1 and abs4-p2 (ATGCAGTCGCCGAATACTGT), and abs4-p3 (TCCTCCGGAGTACCGTCT) and abs4-p4. To determine the copy number of ABS monomers, the read depth of both ABS and the plasmid pACH25 was calculated as the mean of per-base coverage using BEDtools (*52*) and multiplied by two because ABS was heterozygous. The data, with a MAPQ filter of 20, indicated an ABS copy number of 2401, and a pACH25 copy number of 10.5.

The *C2* locus is at 198744424 on chromosome 4. The genetic distance between ABS4 and *C2* is about 12 cM (*53*), as judged by the distance between *C2* and the *tunicate1* gene (which is close to ABS4 at position 183877026).

### Transgenic lines

The *LexA-CENH3(v1*) and *Gal4-CENH3(v1*) constructs were prepared as intronless open reading frames, driven by the maize *CENH3* promoter. The constructs encoded 72 aa from LexA (*54*), based on the crystal structure by (*55*) or 74 aa from Gal4 (*56*). Fused on the C-terminal side of DNA binding domains were 11 aa of maize sequence, 60 aa of oat CenH3 sequence, and the remaining histone fold domain of native maize CENH3. The sequences were codon optimized for maize, commercially synthesized by DNA2.0 (now ATUM, Newark CA), and recombined into pEarleygate vector pEG100 (*57*). The 35S promoter from this vector was then replaced with 1714 bp of the native CENH3 endogenous promoter (using the enzymes BstBI and XhoI) from the inbred HiII (this part of HIII genome is derived from the inbred A188) using the primers 5’-TTCGAAGGCAATTGCAGTAGTGCCT-3’ and 5’-CTCGAGCGCGGTGGGCGCCTCGCA-3’. See Data S1 for full annotated sequences.

The *LexA-CENH3(v2*) construct was based on the *ImmuneCENH3* construct described in (*28*). The guide RNA and ZmPolIII promoter in *ImmuneCENH3* were removed. Then a portion of the gene between the XbaI and BamHI sites was resynthesized by GenScript (www.genscript.com) to include the LexA and oat sequences. The three introns that naturally occur in the sequence that encodes the *CENH3* N-terminal tail were distributed within the LexA and oat sequences at positions where splice site acceptor and donor sequences could be accommodated without altering the encoded amino acid sequence (Data S1). All constructs were transformed into the maize line HiII by the Iowa State University (Ames, Iowa) Plant Transformation Facility (http://www.agron.iastate.edu/ptf/).

The *c2-gfp* allele, identified by (*40*), was ordered as stock tdsgR64H07 from the Maize Genetics Cooperation Stock Center in Urbana Ill. It contains a *trDs\** element in the second exon of the *C2* gene. Fluorescence was viewed using a blue light source and orange filter (Clare Chemical HL34) and imaged using an iPhone. Plants were grown in the UGA Plant Biology greenhouses.

### Chromatin immunoprecipitation and CUT&Tag

Chromatin immunoprecipitation (ChIP) was carried out on individual whole plants as described previously (*58*) using anti-CENH3 (maize) antibodies (*26*). Inputs for each ChIP sample were collected after micrococcal nuclease digestion but before addition of antibodies. Quantitative real-time PCR (qPCR) was carried out in 96-well plates (Bio-Rad #HSP9601) in 15-μL volumes with 2 uL of DNA (0.25 ng/μL) as template, 1 μL primer (5 nM of each, forward and reverse), 4.5 uL water, and 7.5 μL SYBR Select Master Mix for CFX (Applied Biosystems #4472942). Real-time thermocycler conditions were as follows: An initial 50 C for 2 minutes and 95 C for two minutes, then 39 cycles of 95 C for 15 seconds and 60 C for one minute. Primers JIG-261 (5’-CTAGGACTTGCTGGCTTCTATC-3’) and JIG-262 (5’-TCCCTTCTTCGTAAGCTCATTC-3’) amplified the centromeric retrotransposon CRM2, and primers syn3for_short (5’-TCCCTTCTTCGTAAGCTCATTC-3’) and syn3rev_short (5’-TCCGGAGGACAGTCCTCC-3’) amplified ABS. Enrichments were measured as Cq values of each primer pair relative to Cq values from ZmCopia primers (Diagenode #C17040005), which amplified a Copia retrotransposon that is not enriched in centromeres. Each ChIP and input sample was amplified in triplicate for technical replication. For ChIP-seq, a single ChIP and single input library was prepared from each of three plants using a KAPA HyperPrep kit (Roche #KK8502), amplified with 5 cycles of PCR, and Illumina sequenced single-end with 150-nt read lengths. Reads were trimmed of adapters using Cutadapt (*59*) with parameters -q 20 -a AGATCGGAAGAGC -e .05 -O 1 -m 50 and mapped to the Zm-B73-REFERENCE-NAM-5.0 assembly using the Burrows-Wheeler Aligner BWA-MEM with default parameters (*51*). Reads were aligned to CentC, knob180, TR-1, and ABS consensus sequence dimers (*58, 60*) to identify matching reads using blastall with parameters as follows: -p blastn -e 1e-5 -W 7 -G 2 -E 1 -r 1 -q - 1. The ABS sequence is CCAACGAAAATTTCATCCTACTAATTGTGAGCCGCTCACAATTCCAGTAGAAGGCTTACTGTATATATATACAGTATTCGGCGACTGCATCTCTATCACTGATAGGGACTAGCGGTACTCCGGAGGACAGTCCTCCGAGAAGGCATATGTCCCGTTT. For pAHC25 read alignments, BWA-MEM was used with default parameters.

For CUT&Tag, nuclei were extracted from approximately 3 mL finely ground frozen tissue. For leaf samples, this corresponded to about 10 cm length of leaf. For the unfertilized ear, which was 6.4 cm in length, this corresponded to about one-fourth of the ear. A sucrose cushion method of nuclei extraction was used to separate nuclei from abundant amyloplasts (*61*). Rather than diluting the nuclei suspension from the Percoll/sucrose cushion interface in Nucleus Extraction Buffer, the nuclei suspension was diluted in a 1X volume of 1X Wash Buffer from a CUT&Tag-IT Assay Kit (Active Motif 53160). To estimate the yield of nuclei, 50 μl were combined with 50 μl of water and sonicated with a Diagenode Bioruptor for 10 min on high setting with 30-s on-off intervals. DNA concentration was measured using 1 μl of sonicated nuclei with a Qubit dsDNA HS Assay Kit (Invitrogen Q32854). Intact nuclei (not sonicated) containing 150 ng of DNA for leaf or 350 ng for ear was then diluted to a total volume of 1500 μL in 1X Wash Buffer and used as input for the CUT&Tag-IT Assay Kit with both anti-CENH3 (maize) antibodies (*26*) and IgG for each sample (Epicypher 13-0042). Libraries were amplified with 15 or 16 cycles of PCR, Illumina sequenced paired-end 150 nt, and analyzed in the same way as the ChIP-seq libraries (except in paired end mode), and a different adapter sequence was trimmed (CTGTCTCTTATACACATCT).

### Protein extraction and immunoblot analysis

Developing ears approximately 3-4 cm in length were flash-frozen and ground in liquid nitrogen. Total protein was extracted by adding 1 mL of 2X Laemmli buffer (4% SDS, 20% glycerol, 120 mM Tris-Cl, pH 6.8), incubating for 5 minutes at 4°C with gentle rotation, and centrifuging at 15000 x G for five minutes at 4°C. The supernatant was retained and the centrifugation step was repeated. The total protein concentration for each sample was measured using the Qubit Protein Assay Kit (Thermo cat no. Q33211). Approximately 20 μg of each sample were loaded on a 4-20% Mini-PROTEAN TGX Precast Protein Gel (Bio-Rad cat no. 4561094). Proteins were transferred to a Nitrocellulose Membrane (Bio-Rad cat no. 1620145), and the membranes were blocked using TBST containing 5% powdered milk. Rabbit antibodies against maize CENH3 (*26*) or oat CENH3 (*27*) were added to a final dilution of 1:1000 and membranes incubated in primary antibody solution overnight at 4°C with gentle mixing. Membranes were washed three times with TBST and incubated with blocking buffer containing Rabbit IgG HRP Linked Whole Ab (Sigma cat no. GENA934-1ML) at 1:5000 for two hours with gentle mixing. After three final washes in TBST, SuperSignal West Dura Extended Duration Substrate (Thermo Fisher cat no. 34075) was applied to membranes and chemiluminescence was visualized by exposing to X-Ray Film (Research Products International cat no. 248300) in a dark room. Duplicate samples run on the same gels were used for total protein staining with QC Colloidal Coomassie Stain (Bio-Rad cat no. 1610803).

### Fluorescent in situ hybridization

Male meiocytes were prepared for cytological analysis as described in (*62*). Tassels were staged such that anthers containing cells undergoing meiosis were isolated, and anthers were incubated in fixative solution (4% paraformaldehyde, 1% Triton X-100 diluted in PHEMS (30 mM PIPES, 12.5 mM HEPES, 5 mM EGTA, 1 mM MgCl2, 175 mM D-Sorbitol, pH 6.8)) for one hour. Anthers were washed three times in PBS following fixation. Meiocytes were extruded from anthers and immobilized onto polylysine-coated coverslips by centrifuging at 100 x G for one minute then washed three times with PBS. Coverslips containing fixed meiocytes were incubated with 2X SSC (300 mM NaCl, 30 mM sodium citrate, pH 7.0) containing 30% formamide then 2X SSC containing 50% formamide for ten minutes each. Two different sets of labeled oligonucleotides were used for ABS, either four Alexa488-linked oligos (5-TACCGCTAGTCCCTATCAGT-3, 5-CACAATTAGGATGAAAT-3’, 5’-TGTATATATATACAGTAAGC’3’, and 5’-CTTCTCGGAGGACTGTCCTC-3’) or eight Texas Red labeled oligos (5-TACCGCTAGTCCCTATCA-3’, 5’-TGATAGAGATGCAGTCGC-3, 5’-AATACTGTATATATATAC-3’, 5’-TAAGCCTTCTACTGGAAT-3’, 5’-TGAGCGGCTCACAATTAG-3’, 5’-AGGATGAAATTTTCGTTG-3’, 5’-AAACGGGACATATGCCTT-3’, 5’-CGGAGGACTGTCCTCCGG-3’). To detect the Cent4 repeat (*37*), a collection of six Cy5-labeled oligos were used (5’’-ACCCTATGTATCGAAGGA-3’, 5’-CCACTAAAGAACCAAGAT-3’, 5’-CCAAGTAATAGTAAAATA-3’, 5’-GTACAATTTATCCAAACC-3’, 5’-TTAATAAATGTCTAGAGA-3’, 5’-ATGTGATTTGTGTCCAAC-3’). Probe hybridization solution (2X SSC, 50% formamide and each oligo at 1 μM concentration) was administered to meiocytes by suspending coverslips above slides using corners of broken coverslips. Sides of coverslip were sealed with rubber cement, and meiocytes were incubated at 95°C for five minutes, then at room temperature overnight to facilitate probe hybridization. Coverslips were removed from slides and washed for ten minutes each in four subsequent washing solutions: 2X SSC, 20% formamide, 0.01% tween-20; 1X SSC, 10% formamide, 0.001% tween-20; 1X SSC, 1X PBS; 1X PBS. Cells were stained with 0.1 μg/ml DAPI and mounted in 1X PBS containing 100 mg/ml of 1,4 diazobicyclo (2,2,2) octane (DABCO).

Mitotic chromosomes were prepared for cytological analysis as described in (*63*). Roots from freshly germinated seeds were incubated in NO gas for 3.5 hours and then placed on ice. About 1 mm of tissue from the tips of roots were incubated in 2% cellulase Onozuka R-10 (Research Products Internationall) and 1% pectolyase Y-23 (MP Biomedicals) 37°C for 20-50 minutes to digest cell walls. The cells were rinsed twice in 100% ethanol, leaving about ∼4 μL of ethanol in the tube, which was then mixed with 20-30 μL of 100% acetic acid. A dissecting needle was used to break up the tissue and 10 μL of cell suspension was added to slides and allowed to dry. The chromatin was fixed to the slides by UV crosslinking using a total energy of 200 mJ per square cm. 10 μL of probe hybridization solution [5 μL salmon sperm DNA (85 ng/µL salmon sperm DNA in 2x SSC, 1x TE), 4 μL 1X PBS, 0.5 μL of 10 μM Alexa488-labeled ABS oligos, and 0.5 μL of 10 μM Cy5-labeled Cent4 oligos] was added to slides and covered with a plastic coverslip. The slides were incubated at 95°C in a metal incubator for 5 minutes, rinsed with PBS, and mounted with Drop-n-Stain EverBrite Mounting Medium with DAPI (BIOTIUM: 23009). All specimens were imaged using Zeiss Axio Imager.M1 fluorescence microscope with a 63X plan-apo Chromat oil objective. Data were analyzed using Slidebook software (Intelligent Imaging Innovations).

## Data availability

All raw sequencing data generated in this study have been submitted to the NCBI BioProject database (https://www.ncbi.nlm.nih.gov/bioproject/)under accession number PRJNA874319.

## Acknowledgments

We thank the Maize Genetics Cooperation Stock Center for providing maize stocks and Mary Tindall Smith for genotyping stocks. This study was supported in part by resources and technical expertise from the Georgia Advanced Computing Resource Center, a partnership between the University of Georgia’s Office of the Vice President for Research and Office of the Vice President for Information Technology. This work was supported by grants from the National Science Foundation (IOS-1444514 and IOS-2040218).

