## Supplemental Figures and Data for "Synthetic maize centromeres transmit chromosomes across generations"

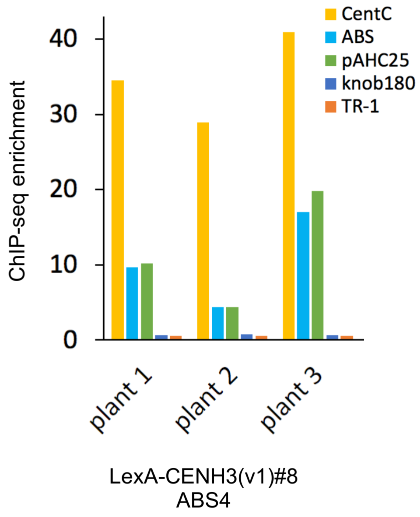

**Figure S1.** Confirmation that LexA-CENH3(v1) recruits native CENH3 by a ChIP-seq assay. The data show CENH3 ChIP enrichment relative to input, measured by Illumina sequencing. Each ChIP is from an individual plant that was heterozygous for ABS4 and the *LexA-CENH3(v1)* transgene. CentC is a native centromere repeat whereas knob180 and TR-1 are non-centromeric repeats. The Y-axis indicates the numbers of reads in the ChIP sample divided by numbers in input and normalized by the total number of reads.

| Cross | Transgene genotypes | *cenh3* genotypes |
|---|---|---|
| *-/LexA-CENH3(v1)*<br>*+/cenh3*<br>⊗ | *-/LexA-CENH3(v1) or*<br>*LexA-CENH3(v1)/LexA-CENH3(v1)* (**67**) | +/+ (33) |
| | | +/*cenh3* (34) |
| | | *cenh3/cenh3* (**0**) |
| | -/- (11) | |
| *-/LexA-CENH3(v2)*<br>*+/cenh3*<br>⊗ | *-/LexA-CENH3(v2) or*<br>*LexA-CENH3(v2)/LexA-CENH3(v2)* (**20**) | +/+ (7) |
| | | +/*cenh3* (9) |
| | | *cenh3/cenh3* (**4**) |
| | -/- (4) | |

**Figure S2**. Complementation of the *cenh3* null by LexA-CenH3 transgenes. A minus sign indicates the lack of *LexA-CENH3* transgene. A plus sign indicates the wild-type *CENH3* allele. Plants heterozygous for either *LexA-CENH3(v1)* or *LexA-CENH3(v2)* were crossed to lines carrying the *cenh3* null and heterozygotes self-crossed. The genotypes of progeny are shown to the right. Note that of 67 plants carrying *LexA-CENH3(v1)*, none were homozygous for cenh3, indicating that the transgene cannot complement the null. In contrast, of 20 plants carrying *LexA-CENH3(v2),* four were homozygous for cenh3. One of these four plants, which proved to be homozygous for *LexA-CENH3(v2)*, looked normal (see Figure S5).

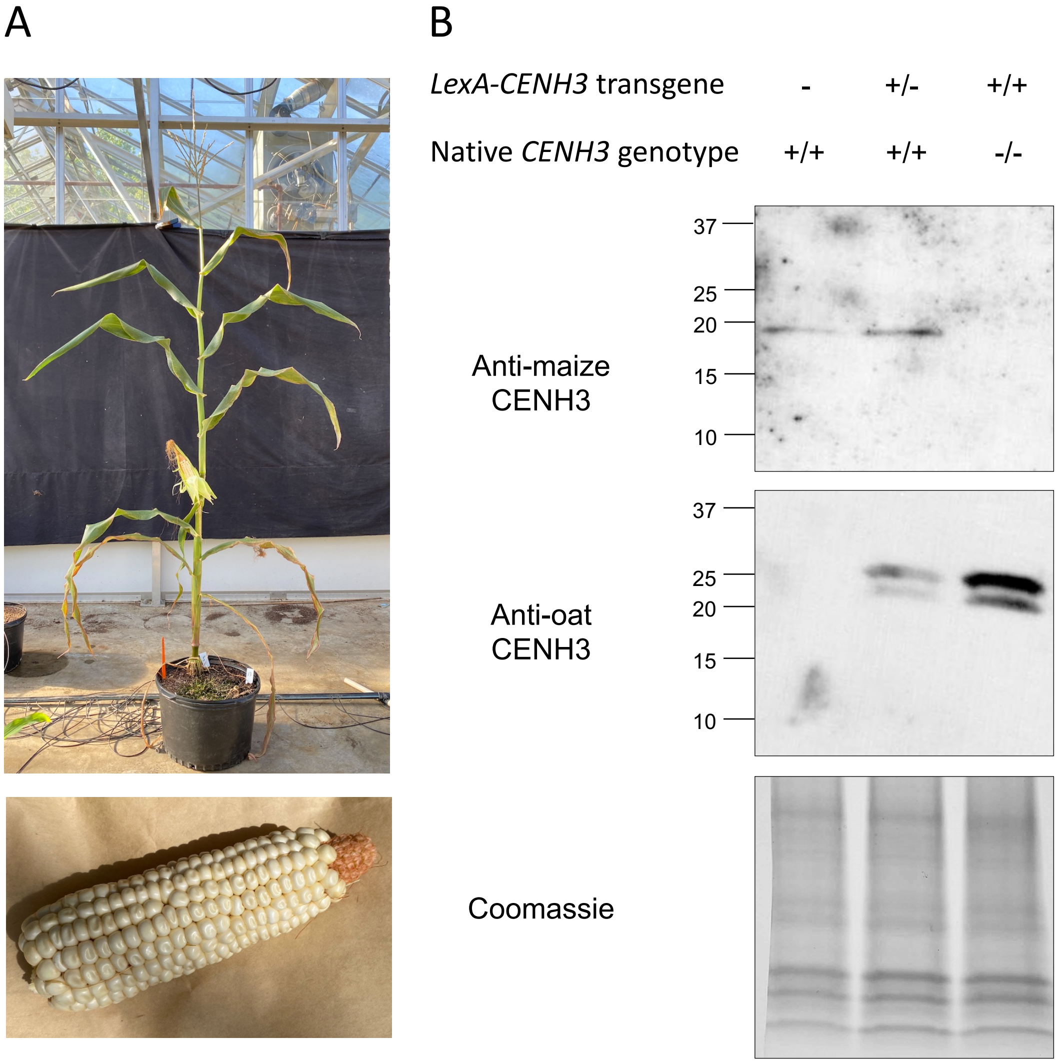

**Figure S3**. Demonstration that *LexA-Cenh3(v2)* is sufficient for cell division and plant growth. **A**). A plant homozygous for the *LexA-CENH3(v2)* transgene and homozygous for a *cenh3* null mutant. The plant was self-crossed to reveal a full ear (below), indicating that the transgene was homozygous (one quarter of the seeds would have been dead if the plant was heterozygous for *LexA-Cenh3(v2)*). The second ear of this plant was used for the third lane of the protein blot in B. **B**) Protein blot analysis of CENH3 in plants of different genotypes. A minus sign indicates the lack of *LexA-CENH3* transgene. A plus sign indicates wild-type *CENH3* allele. Native maize CENH3 is 17 kDa and recognized by a maize-specific antibody. The oat CENH3 antibody recognizes the LexA-CENH3 fusion protein only. The predicted ~27 kDa LexA-CENH3 band is observed along with a smaller band of unknown cause/origin. Importantly, native CENH3 was not detectable in the *LexA-CENH3(v2)/LexA-CENH3(v2), cenh3/cenh3* null plant (third lane). The lower panel (Coomassie) shows that the amounts of protein loaded into each lane were similar.

A

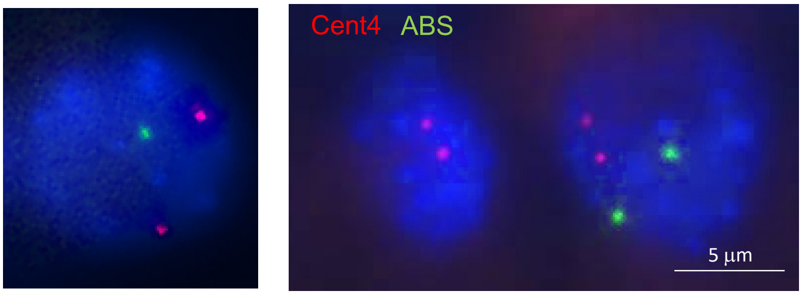

| | 1 ABS | 0 ABS | 2 ABS |
|---|---|---|---|
| Plant #1 | 56 | 8 | 6 |
| Plant #2 | 97 | 3 | 8 |
| Plant #3 | 138 | 5 | 3 |
| Plant #4 | 166 | 5 | 11 |
| Plant #5 | 84 | 3 | 5 |
| Total | 541 | 24 (4.0%) | 33 (5.5%) |

B

| | No recombination 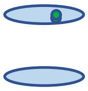 | Yes recombination but no bridge 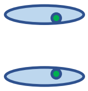 | Presumed recombination, bridge present and ABS staining in midzone 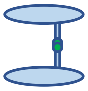 |
|---|---|---|---|
| Plant #2 | 17 | 10 | 17 |
| Plant #12 | 5 | 3 | 2 |
| Plant #16 | 8 | 1 | 5 |
| Total | 30 | 14 | 24 |

**Figure S4**. Segregation errors in *LexA-CENH3(v2)* ABS4 plants. **A**) Mitotic errors. Root tips from five sectored seeds were processed for FISH and the number of Cent4 and ABS spots counted in interphase nuclei. Two Cent4 (red) spots were observed in all nuclei. ABS (green), which is on one copy of chromosome 4, showed high frequencies of mis-segregation consistent with the model in Figure 2A. The images shown above are examples. The two cells in the right image are daughter cells of a single division, where the left cell has no ABS and the right cell as two ABS loci. **B**) Meiotic errors. Three plants heterozygous for *LexA-CENH3(v2)* and ABS4 were analyzed at meiosis (siblings from a single ear). Only cells in mid-anaphase I, identified by the short distance between segregating chromosomes masses, were tallied. Figures with no recombination between the centromere and ABS4 were identified by having ABS dots only on one side (first column). Cases where recombination occurred and ABS4 loci segregated freely to one pole were identified by having one ABS dot on both sides (second column). Figures with bridges invariably had one or two ABS dots in the midzone, suggesting there had been recombination between centromere and ABS4, and that centromere cohesion at ABS4 restrained movement. At late anaphase I and telophase I, no intact bridges were observed.

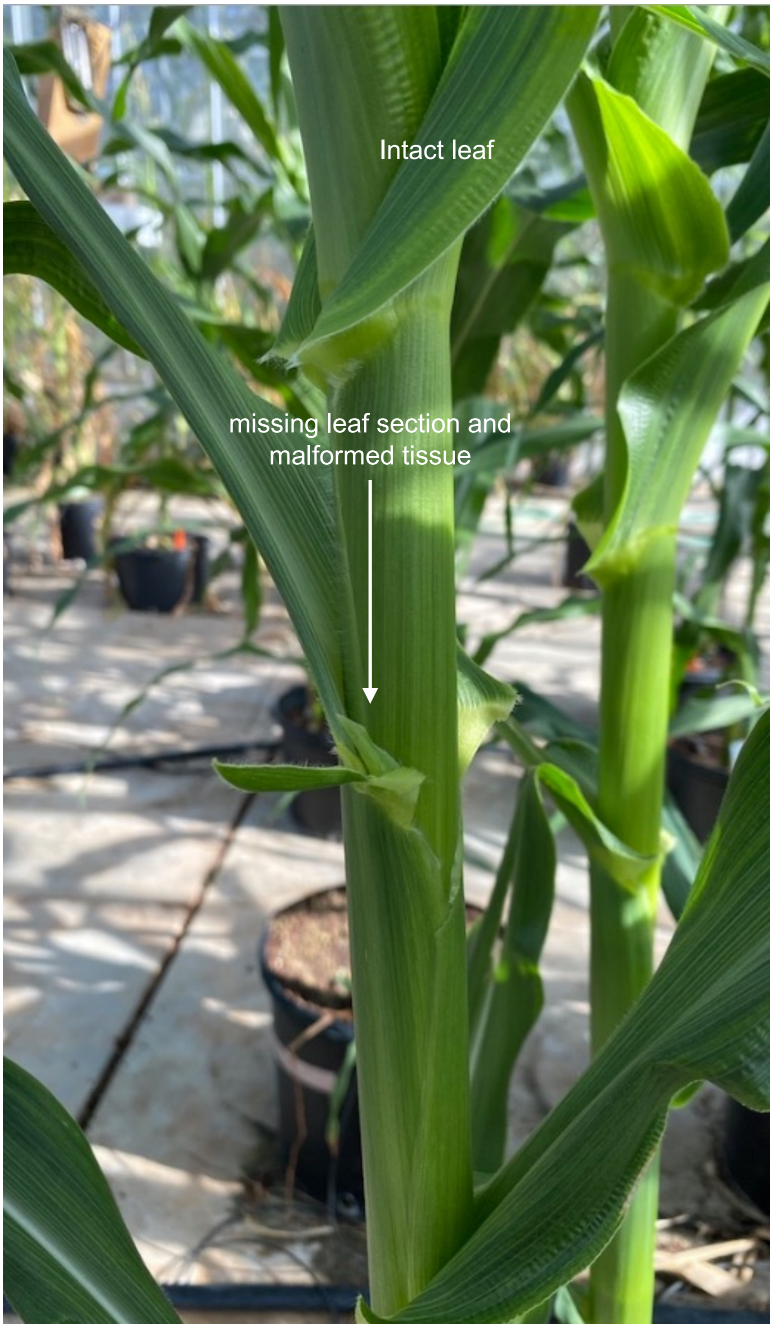

**Figure S5**. Leaf defects observed in plants with both *LexA-CENH3(v2)* and ABS4. The leaves of nearly all *LexA-CENH3(v2)* ABS4 plants looked and felt uneven, with ridges or crinkles. Occasionally the leaves had missing pieces, such as shown here.

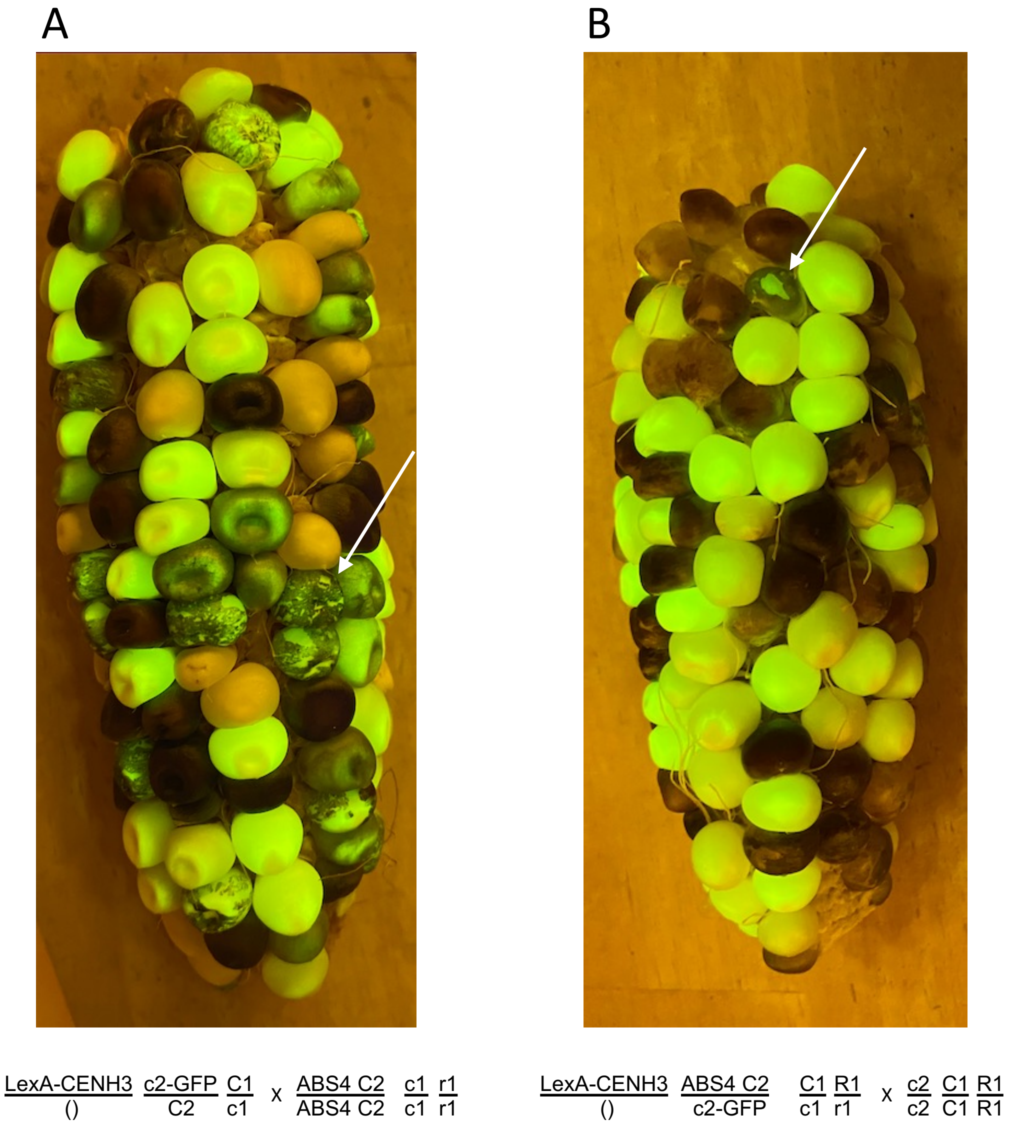

**Figure S6**. Crosses involving c2-GFP. **A**). The cross used to generate material for the neochromosome screen. *C2* confers purple pigmentation to the outer layers of the endosperm (purple appears black under the blue light used here). The binding of LexA-CENH3 to ABS4 causes errors in chromosome segregation and loss of the linked *C2* gene in sectors (arrow). Two other recessive color alleles (*c1* and *r1*) are also segregating in these transgenic lines, which caused many kernels to be colorless regardless of the presence of *C2* (in this case only *c1* was heterozygous on the female side). **B**) Fluorescent sectored kernels from A were planted and crossed as females to a *c2* tester. Kernels that received both *C2* (purple aleurone) and *c2-GFP* (fluorescent endosperm) were candidates for having inherited a neochromosome. This ear shows such a kernel. Filing off the outer layer of cells removes the purple pigment and makes the underlying fluorescence more obvious (arrow).

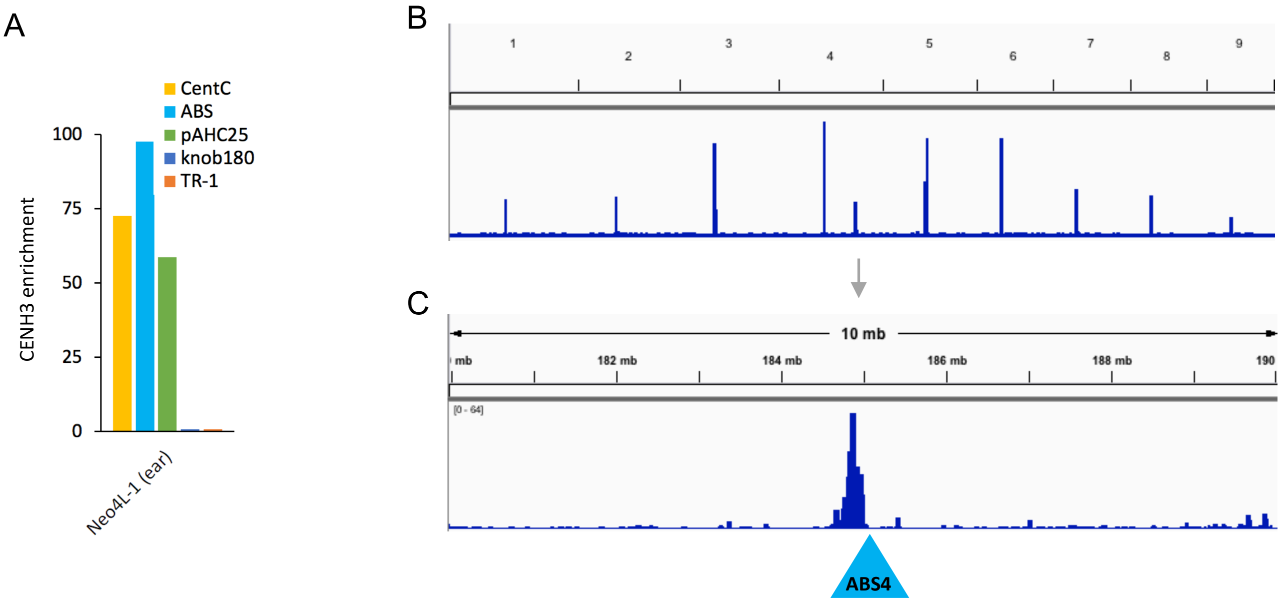

**Figure S7**. Analysis of CENH3 CUT&Tag from a developing ear carrying Neo4L-1. **A)** CENH3 enrichment relative to IgG of repetitive DNA elements. CentC is a native centromere repeat, knob180 and TR-1 are non-centromeric repeats. **B)** CENH3 peaks on all ten chromosomes. **C**) CENH3 peak over a 10-MB window from chromosome 4. Blue triangle indicates ABS4 insertion site in B73 v5 genome coordinates. A peak of enrichment is observed from ~184650000 to 185000000 (at 50-Kb level resolution).

### Supplemental Tables

**Table S1.** Results of the purple/fluorescent genetic screen for potential neochromosomes

| | | | | | |
|---|---|---|---|---|---|
| **Fall 2021** | # kernels in each class | | | | |
| Cross | *c2-gfp/c2* | *C2/c2* | *C2/c2-gfp/c2* | | |
| KD4210-4 X 4211 | 199 | 166 | **1** | → **Spring 2022**: | Potential translocation |
| KD4210-5 X 4211 | 128 | 87 | 0 | | |
| KD4210-2 X 4211 | 135 | 110 | **2** | → **Spring 2022**: | 1 did not have ABS |
| | | | | | 1 contained Neo4L-2 |
| **Spring 2022** | # kernels in each class | | | | |
| Cross | *c2-gfp/c2* | *C2/c2* | *C2/c2-gfp/c2* | | |
| KD4231-1 X 4245 | 52 | 34 | **9** | → **Summer 2022**: | All did not have ABS |
| KD4231-2 x 4244 | 140 | 35 | 2 | | |
| KD4231-3 x 4244 | 146 | 12 | 1 | | |
| KD4231-4 x 4244 | 139 | 86 | 5 | | |
| KD4231-5 x 4244 | 115 | 8 | 0 | | |
| KD4231-6 x 4244 | 174 | 21 | 1 | | |
| KD4231-7 x 4244 | 161 | 50 | 4 | | |
| KD4231-8 x 4244 | 170 | 79 | 3 | | |
| KD4231-9 x 4244 | 39 | 3 | 0 | | |
| KD4231-10 x 4244 | 229 | 146 | 5 | | |
| KD4231-11 x 4244 | 156 | 113 | 1 | | |
| KD4231-12 x 4244 | 149 | 44 | 3 | | |
| KD4231-13 x 4244 | 188 | 104 | 7 | | |
| KD4231-14 X 4244 | 345 | 194 | 4 | | |
| KD4232-1 X 4245 | 131 | 74 | **5** | → **Summer 2022**: | 2 did not have ABS |
| KD4232-2 x 4245 | 43 | 26 | 1 | | 2 yielded no seeds when crossed |
| KD4232-3 x 4244 | 146 | 100 | 0 | | 1 contained Neo4L-3 |
| KD4232-4 X 4244 | 169 | 89 | 0 | | |
| KD4232-5 x 4244 | 135 | 59 | 2 | | |
| KD4232-6 x 4244 | 144 | 25 | 0 | | |
| KD4232-7 x 4244 | 260 | 1 | 0 | | |
| KD4232-8 x 4244 | 195 | 128 | 4 | | |
| KD4232-9 x 4245 | 51 | 37 | 2 | | |
| KD4233-1 X 4245 | 35 | 12 | 0 | | |
| KD4234-1 x 4245 | 102 | 0 | 0 | | |
| KD4234-2 x 4244 | 167 | 13 | 1 | | |
| KD4234-3 x 4244 | 271 | 98 | 6 | | |
| KD4234-4 x 4244 | 124 | 13 | 5 | | |
| KD4234-5 x 4244 | 223 | 115 | 2 | | |
| KD4234-6 x 4244 | 262 | 54 | 0 | | |
| KD4234-7 X 4245 | 271 | 251 | 2 | | |
| KD4235-1 x 4244 | 89 | 64 | 3 | | |
| KD4235-2 X 4244 | 68 | 8 | 0 | | |
| KD4235-3 x 4244 | 154 | 83 | 1 | | |
| KD4235-4 x 4244 | 219 | 68 | 2 | | |
| KD4236-1 x 4245 | 209 | 38 | 6 | | |
| KD4236-2 x 4245 | 146 | 12 | 0 | | |
| KD4236-3 x 4244 | 62 | 24 | 0 | | |
| KD4236-4 x 4244 | 73 | 36 | 1 | | |
| KD4236-5 X 4245 | 197 | 62 | 0 | | |
| KD4236-6 X 4245 | 149 | 51 | 4 | | |
| KD4236-7 x 4244 | 168 | 89 | 6 | | |
| KD4236-8 x 4245 | 78 | 14 | 2 | | |
| KD4237-1 x 4245 | 205 | 101 | 3 | | |
| KD4237-2 x 4245 | 135 | 14 | 2 | | |
| KD4237-3 X 4245 | 120 | 5 | 1 | | |
| KD4237-4 X 4244 | 156 | 47 | 0 | | |
| KD4237-5 X 4245 | 63 | 33 | 0 | | |
| KD4237-6 X 4245 | 166 | 127 | 4 | | |
| KD4238-1 X 4244 | 126 | 16 | **6** | → **Summer 2022**: | 5 did not have ABS |
| KD4238-2 x 4245 | 130 | 9 | 1 | | 1 contained Neo4L-4 |
| KD4238-3 x 4245 | 58 | 14 | 0 | | |
| KD4238-4 x 4244 | 26 | 16 | 0 | | |
| KD4238-5 X 4245 | 79 | 30 | 1 | | |
| KD4238-6 x 4245 | 157 | 21 | 3 | | |
| KD4238-7 x 4245 | 95 | 47 | 2 | | |
| KD4238-8 x 4245 | 217 | 125 | 7 | | |
| KD4239-1 x 4245 | 164 | 95 | 3 | | |
| KD4239-2 X 4245 | 97 | 22 | 1 | | |
| KD4239-3 x 4244 | 110 | 42 | 1 | | |
| KD4239-4 x 4244 | 118 | 4 | 1 | | |
| KD4239-5 x 4244 | 126 | 37 | 4 | | |
| KD4241-1 X 4245 | 40 | 36 | 0 | | |
| KD4241-2 X 4244 | 164 | 112 | 1 | | |
| KD4242-1 X 4244 | 268 | 133 | 2 | | |
| KD4242-2 X 4244 | 95 | 43 | 1 | | |
| KD4242-3 X 4245 | 114 | 79 | 0 | | |
| KD4242-4 x 4244 | 166 | 6 | 8 | | |
| KD4242-5 x 4244 | 259 | 156 | 5 | | |
| KD4242-6 x 4244 | 327 | 176 | 4 | | |
| KD4242-7 x 4244 | 222 | 183 | 0 | | |
| KD4243-1 X 4244 | 218 | 29 | 2 | | |
| KD4243-2 x 4244 | 279 | 11 | 5 | | |
| KD4246-1 x 4244 | 185 | 44 | 6 | | |
| KD4246-2 x 4244 | 126 | 10 | 4 | | |
| | 11817 | 4759 | 181 | 16757 total | 1.09% purple and fluorescent |

**Table S2**. Segregation data for Neo4L chromosomes

| | Crossed as female | C2 | total | % C2 | # plants from C2 seed that did not have ABS[1] | | Crossed as male | C2 | total | % C2 | # C2 seed that were sectored | # plants from C2 seed that did not have ABS[2] |
|---|---|---|---|---|---|---|---|---|---|---|---|---|
| **Neo4L-1** | | | | | | | | | | | | |
| 1st gen | KD4205-5 X 4202 | 26 | 237 | **10%** | | | KD4202 X 4205-5 | 10 | 140 | **7%** | | 7/10 |
| 2nd gen | KD4212-6 X 4211 | 33 | 93 | **26%** | 2/8 | | KD4211 X 4212-6 | 41 | 184 | **22%** | 10/41 | |
| 3rd gen | KD4246-4 X 4244 | 70 | 266 | **21%** | 1/9 | | KD4245 X 4246-4 | 29 | 180 | **16%** | 6/29 | |
| | KD4246-5 X 4244[3] | 60 | 240 | **20%** | 3/10 | | KD4245 X 4246-5 | 22 | 189 | **12%** | 1/22 | 5/10 |
| | KD4246-6 X self [4] | 135 | 219 | 62% | | | KD4244 X 4246-6 | 73 | 348 | 21% | 12/73 | 9/18 |
| 4th gen | KD4301-3 X 4277 | 49 | 182 | **21%** | 1/9 | | KD4277 X 4301-3 | 21 | 266 | **7%** | 4/21 | 8/10 |
| | KD4302-3 X 4277 | 56 | 207 | **27%** | | | KD4277 X 4302-3 | 3 | 51 | **6%** | | |
| | KD4302-4 X 4277 | 62 | 199 | **31%** | | | KD4277 X 4302-4 | 5 | 41 | **12%** | | |
| **Neo4L-2** | | | | | | | | | | | | |
| 1st gen[5] | KD4249-1 X 4244 | 59 | 214 | **22%** | 0/24 | | KD4245 X 4249-1 | 0 | 214 | **0%** | | |
| 2nd gen[6] | KD4300-4 X 4277 | 48 | 171 | **28%** | | | N/A[7] | | | | | |
| | KD4300-5 X 4277 | 85 | 320 | **27%** | 4/18 | | N/A | | | | | |
| | KD4300-7 X 4277[8] | 61 | 230 | **27%** | 3/20 | | KD4277 X 4300-7 | 3 | 35 | **9%** | 0/3 | |
| | KD4300-8 X 4277 | 49 | 200 | **25%** | | | N/A | | | | | |
| **Neo4L-3** | | | | | | | | | | | | |
| 1st gen | KD4303-2 X 4277 | 31 | 124 | **25%** | 2/6 | | KD4277 X 4303-2 | 53 | 392 | **14%** | 5/53 | 6/10 |
| **Neo4L-4** | | | | | | | | | | | | |
| 1st gen | KD4304-1 X 4277 | 65 | 238 | **21%** | 0/9 | | KD4277 X 4304-1 | 22 | 212 | **10%** | 6/22 | 5/10 |

[1]Among progeny from female crosses, 16/113 (14%) of the seeds with C2 endosperm gave rise to seedlings that did not have ABS

[2]Among progeny from male crosses, 40/68 (59%) of the seeds with C2 endosperm gave rise to seedlings that did not have ABS

[3]48 colorless seeds were planted and 1 had ABS.

[4]62% C2 on self cross is not consistent with ~20% transmission through male & female. The plant may have had two copies of Neo4L-1.

[5]This plant carried LexA-CENH3(v2).

[6]These plants lacked LexA-CENH3(v2).

[7]All plants carrying Neo4L-2 in this generation exerted few anthers and had little pollen.

[8]39 colorless seeds were planted and none carried ABS.

**Supplemental Data**

**Data S1.** Illustrations and sequences of *LexA-CENH3(v1)* and *LexA-CENH3(v2)*.

**Comparison of LexA-CENH3(v1) and LexA-CENH3(v2)**

LexA-Cenh3(v1)
3300 bp

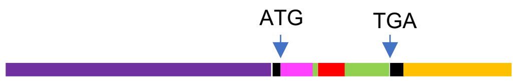

Promoter
Maize mRNA
LexA
Oat
Vector
OCS terminator

LexA-Cenh3(v2)
7027 bp

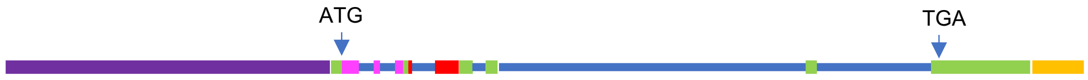

Promoter
Maize mRNA
LexA
Oat
Intron
PinII terminator

**LexA-CENH3(v1) transgene DNA sequence**

Promoter from A188, Vector sequence, LexA, Maize mRNA , oat, OCS terminator

AGGCAACTGCAGTAGTGCCTCTGTTTTAGAGTGTAACTACAGATTTGTCCCTATTTTTTTAGAGTTTGCGTGTTTGT
CCCTATTTTTTTTTCAAATCAAACTATTGTATACCCCTACTCCATTAGTTATACTTAACGGTGTTAACTCTTGATAA
AAAGACAAGGGATAATTAGATTAGTGCCACTGTATTAGAGTGTAATGATAGTTTTGACCGATGTTTTAGACTTCACA
TGTTTTTATGACAATTCAAATTGTTTCCATAACATTTTAAATTATTTTGACAACATTTAGAATTGTTTTGCAATAAT
TTAAATTATTTCCAAAATAAAAATATTTTGACAATTATTTTATCAACAAATTATTTTTTTACAAAATAATTTGTCAA
GATACTTTTTTTAATTTTTTTTGAAAATAATCAAATTATTGTAAATATAATTTGAATTGTTGCAAAAATAATTTAGA
TTGTCATAAAAACACGTAAGTCTAAAAATAGGCGTAAAACTACAGTTATACTCTAAAAAAGTGGTAATGGTGTAATT
GTTCCTTGTCTATTTTATCAAGACATAGCACCGTTAAGTACAACTAACGGAGTAGGGATATACAATATTTTTGTTTT
AAAAAATAGGGGTAAATACGCAAAATCTAAAAACAGAGGTAAAACTACAATTACATGTTAAAATAGGGGCATCAGTA
CAATTGTATCTTTTATATAGCAGCATGCGCCCTGTTGGGATACAACCTGAGCCTTCACATGTCTTCTAGATGGTTCC
CAACCCTTTGGCCAAGATCGTACGGATAATATTGCGAGGAACCCAAATCAACGGTGTCTATGTTATGTTGATGTGGA
TGGTTTACCTGGGCGCAAAAGTGCGCTTGTTTCGTCTGTACAAATATACTTTAAGTATGGTTTTGATCTTTTTATTT
TTCATTTTTTAAATAAACGAGCCAATCAAACCTGATATAAAAAATCAAATGAATTATAAATAGAGACGGAGAGAGTA
TATATATTTGTTTTACTATTGTTTAAAGTATTAAAATATAGTGGACGGAGGAACGTTCTCTATGTTTAAAAGGACGT
TTTAGAGGACGTTGTGTTGTTGAAGGAAATATGAAAAAAATCTTCTGCATATTTAGAAGGGAGGAGCGTTTACACAT
TACTTTCGGGACTTCAACCCAAATATGTCAAGGTTTGTGAGTGGCTCAGTGCAGAAAAAAATCATGTATATACCAAA
TGTAAATAATATCTTTTACAACCCATCACATTCACATTTGGAGGTTCACAAAGATAGATCAAAATTTATAAAATAAT
CCTTTAATAATTTTTTCTTCTTCTATTTATATGGACCATTAGCTGGTGTATATGAGGAGCTGTAAAAGATATTCTTT
ACACCAGAGATGTAAAGAATTTTTTTAACTCAATGCTGGTTACCGGCTGGGAGGACGATGATAAAGAAAGCATCTCT
CACTGCATTCCGGGCCCACTACTCAAACGTTCGGCACGCCAGGTTGGCAGGTAGCCGTTACCTCGATAGGCACTCGG
CCACTCGCACGCAGACACCACACCAGTGTGCTCAGTGCTCACTGCTCACCATAACGCTGCACCTCTTTTCATTTCAC
CATCTCCTGCCCCCTTAAAAAAAAAACTCACCGTCGACACGCCCTCCCGTCCCGAGAGTTCTGAATCGAAACCGTCG
GCCACGAGAGCAGTGCGAAGCGCCCACCGCGCTCGAGATCACAAGTTTGTACAAAAAAGCAGGCTCCGCGGCCGCCC
CCTTCACCATGAAGGCCCTGACCGCCAGGCAGCAAGAGGTGTTCGACCTGATCAGGGACCACATCTCCCAGACCGGC
ATGCCGCCGACCAGGGCCGAGATCGCCCAGAGGCTGGGCTTCAGGTCCCCGAACGCCGCGGAGGAGCACCTGAAGGC
CCTGGCCAGGAAGGGCGTGATCGAGATCGTGTCCGGCGCCTCCAGGGGCATCAGGCTGCTGCAGGAGGAGATGGCTC
GAACCAAGCACCAGGCCGTGAGGAAGTCGAAGCCGACGCCCAAGAAGCAGCTCAAGTTCGGCCGCTCCCCGGGCCAG
ACGGCGGAGCAGGAGACAGGCGGCCCGAGCACGTCGGCGGCACCAAGGCGAGGCGCGCGGAGGCCAGCAGCAACGCC
GGCTCCAGGGGCACCTGCGCAACAGAGGGCGAGGAAGCCGCATCGGTTCAAGCCAGGGACTGTAGCGCTGCGGGAGA
TCAGGAAGTACCAGAAGTCCACTGAACCGCTCATCCCCTTTGCGCCTTTCGTCCGTGTGGTGAGGGAGTTAACCAAT
TTCGTAACAAACGGGAAAGTAGAGCGCTATACCGCAGAAGCCCTCCTTGCGCTGCAAGAGGCAGCAGAATTCCACTT
GATAGAACTGTTTGAAATGGCGAATCTGTGTGCCATCCATGCCAAGCGTGTCACAATCATGCAAAAGGACATACAAC
TTGCAAGGCGTATCGGAGGAAGGCGTTGGCATGAAAGGGTGGGCGCGCCGACCCAGCTTTCTTGTACAAAGTGGTG
CCTAGGTGAGTCTAGAGAGTTAATTAAGACCCGGGACTAGTCCCTAGAGTCCTGCTTTAATGAGATATGCGAGACGC
CTATGATCGCATGATATTTGCTTTCAATTCTGTTGTGCACGTTGTAAAAAACCTGAGCATGTGTAGCTCAGATCCTT
ACCGCCGGTTTCGGTTCATTCTAATGAATATATCACCCGTTACTATCGTATTTTTATGAATAATATTCTCCGTTCAA
TTTACTGATTGTACCCTACTACTTATATGTACAATATTAAAATGAAAACAATATATTGTGCTGAATAGGTTTATAGC
GACATCTATGATAGAGCGCCACAATAACAAACAATTGCGTTTTATTATTACAAATCCAATTTTAAAAAAAGCGGCAG
AACCGGTCAAACCTAAAAGACTGATTACATAAATCTTATTCAAATTTCAAAAGTGCCCCAGGGGCTAGTATCTACGA
CACACCGAGCGGCGAACTAATAACGCTCACTGAAGGGAACTCCGGTTCCCCGCCGGCGCGCATGGGTGAGATTCCTT
GAAGTTGAGTATTGGCCGTCCGCTCTACCGAAAGTTACGGGCACCATTCAACCCGGTCCAGCACGGCGGCCGGGTAA
CCGACTTGCTGCCCCGAGAATTATGCAGCATTTTTTTGGTGTATGTGGGCCCCAAATGAAGTGCAGGTCAAACCTTG
ACAGTGACGACAAATCGTTGGGCGGGTCCAGGGCGAATTTTGCGACAACATGTCGAGGCTCAGCAG

**Gal4-CENH3 transgene DNA sequence**

Same as LexA-CENH3(v1), except the LexA sequence was replaced with:
ATGAAGCTGCTCTCCTCTATCGAGCAGGCCTGCGACATCTGCAGGCTGAAGAAGCTGAAGTGCTCCAAGGAGAAGCC
GAAGTGCGCCAAGTGCCTGAAGAACAACTGGGAGTGCAGGTACTCCCCGAAGACCAAGAGGTCCCCGCTGACCAGGG
CCCACCTGACCGAGGTGGAGTCCAGGCTGGAGAGGCTGGAGCAGCTGTTCCTGCTCATCTTCCCGAGG

**LexA-CENH3(v2) transgene DNA sequence**

Promoter, LexA, Maize mRNA, oat, PinII terminator

GGCGCGCCGAGCAGGGCGAAAAGAAAGATCAAGTACGACTTGTTAAACTTGACAACTCACCAACATTGTCATCATTG
GTTAAGCCATTTTCGAGACCCAACAATGAGACATGGATAAAGATGGATCAAGATTATACTAACAATGAGCTAATGGA
GGAGCATAATGGCCTGATCATAAATGAATGAGCAATGCAGGGAACCATGCTTGTTTTTTTGTAATCTATATGCAAAT
GTTTCGTCAAGGATTTGGTCTATACCTCTTTTTCCTTGCTGCACTCCTAGATCACCATAAGTGAGGGCTCTCCTTGC
AATGTGACAACTAACCATAGCCATGATTCTTCCTCGTATTTTTTTACTTTTGTTTTTTCTTTCTCCCTCTCTTTTAC
ATGCGTTCTCCTTTGGTCTCTAGTGTTTATCTGGCATACTAGTATCTTAGTCACTTCTCTCTTTTTATGTAGAGGCA
ATTGCAGTAGTGCCTCTGTTTTAGAGTGTAACTACAGATTTGTCCCTATTTTTTTAGAGTTTGCGTGTTTGTCCCTG
TTTTTTCAAATCAAACTATTGTATACCCCTACTCCATTAGTTATACTTAACAATGTTAAGTCTTGATAAAAAGACAA
GGGATAATTGGATTAGTGACCCTGTTTTAGAGTGTAATTATAGCTTTGCCCGATGTTTTAGACTTCACATGTTTTTA
TGACAATTCAAATTGTTTCCATAACATCTTAAATTATTTTGACAACATTTAGAATTGTTTTGCAATAATTTAAATTA
TTTCCAAAATAAAAATATTTTGACAATTATTTTATCAACAAATTAAATTATTTTTTTTACAAAATAATTTGTCAAGG
TACTTTTTTAAAATTTTGAAAATAATCAAATTATTGTAAATATAATTTGAATTGTTGCGAAAATAATTTGGATTGTC
ATAAAAACACGTAAGTCTAAAAATTAGGCGTAAAACTACAATTATACTCTAAAAAGGTGGTAATGGCGTAGTTGTTC
CTTGTCTATTTTATCAAGACATAGCACCGTGCAGTACAACTAATGGAGTAGTGACATACAACAATTTTGTTTTAAAT
AATAGGGGTAAATACGCAAAGTCTAAAAAACAGAGGTAAATCTACAATTACATGTTAAAATAGAGGCATCGATACAA
TTGTACCTTTTATATAGCAGCATGCGCCCTGTTGGGATACAATTGTACCTTTCACATGTCTTCTAGATGGTTCCCAA
CCCTTTGGCCAAGATCGTACAGATAATATTGCGAGGAGCCCAAATCAACGGTGTCCATATGTTATGTTGATGTGGAT
GGTTTACCTAGGCGCAAAAGTGCGCTGGTTTCGTCCGTACAAATATACTTTAAGTATGGTTTTGATTTTTTTCTATT
TTTCATTTTTTAAATAAAACGAGACAATCAAATCTGATATAAAAATCAAATGAATTATAAATAGAGACGGAAAGAGT
ATATATATTTGTTTTGCTATTATTTAAAGTATTAAAAGATAGTGGACGAATGAACGTCCTCTATGTTTAAAAGAACG
TTTTAGAGGACGTTGTGTTGTTGAAGGAAATATGAAAAAAAAATCTTCTGCATATTTAGAAGGGAGGAGCGTTTACA
CATTACTTTCGGGACTTCAACCCAAATATGTCAAGGTTTGTGAGTGGCTCAGTGCGGAAAAAAAATCCTATATATAC
CAGATGTAAACACTATCTTTTACAGCCTATCACATTCACATTTAGAGGTTCACAAAGATAGATCAAAATTTATAAAA
TAATCATTTAATATTTTTTTTATTTTATTTATATGGATAAGCAGCTGGTGTATGTGAGGAGCTGTAAAAGATATTTT
TTACATCCGAGATGTAAAGATTTTTTTTAACTCAATGCTGGTTACCGGCTGGGAGGACGATGATAAAGAAAGCATCT
CTCACTGCATTCCGGGCCCACTACTCAAACGTTCGGCACGCCAGGTTGGCAGGTAGCCGTTACATCGATAGGCACTC
GGCCACTCGCACGCAGACACCACACCAGTGTGCTCAGTGCTCACTGCTCACCATAATAACGCTGCACCTCTTTTCAT
TTCACCATCTCCTGCCCCCTTAAAAAAAAGACTCACCGTCGACACGCCCTCCCGTCCCGAGAGTTCTGAATCGAAAC
CGTCGGCCACGAGAGCAGTGCGAGGCGCCCACCGCGATGAAGGCCCTGACCGCCAGGCAGCAAGAGGTGTTCGACCT
GATCAGGGACCACATCTCCCAGACCGGCATGCCGCCGACCAGGGCCGAGATCGCCCAGAGGCTGGGCTTCAGGTAAC
CCGGGTCCCGCGCTCCCCCCCGCTTCGCAAGCAGACGCTGTCGCTTCTCTCCGACCCTGGTGCTAAGCACGTTCCTT
GTTCCGTCTTTTGCAGGTCCCCGAACGCCGCGGAGGAGCACCTGAAGGCCCTGGCCCGGTGAGCGCGTGCGTGCGGG
GATCAGTTCCCTCCTTTTGCCTTTTTTTGTTGGGCTGCTCTTACTTGCTTGCAAGCTGTTTGATGGAATGCAGGAAG
GGCGTGATCGAGATCGTGTCCGGCGCCTCCAGGGGCATCAGGCTGCTGCAGGAGGAGATGGCTCGAACCAAGCACCA
GGCCGTGAGGAAGTCGAAGCCGACGCCCAAGAAGCAGGTGAGGACCTATTTGTCGTTGCTGGATGCTGGGTTTCGCT
TGCAATCTAATTTTGTTGCAAGATGAGGGCGAATGTGCCAGTTCCATGTGGGTGTCATGGTCTCGGAGTTACTACCT
TAATTGCTCACCATAGTATGTTTTCTTAAAAAAAACAGTTGAAGTTCGGCCGCTCCCCGGGCCAGACGGCGGAGCAG
GAGACAGGCGGCCCGAGCACGTCGGCGGCACCAAGGCGAGGCGCGCGGAGGCCAGCAGCAACGCCGGCTCCAGGGGC
ACCTGCGCAACAGAGGGCGAGGAAGCCGCATCGGTTCAAGCCAGGTACGGTCGCCCTGCGCGAGATCAGGAAGTACC
AGAAGTCCACTGAACCGCTCATCCCCTTTGCGCCTTTCGTCCGTGTGGTGGGTGCAGGCGTGTTTGTCCTCTGCATA
GTATGGGGTTGTTCCGCATTCTGTCTAATGGAAAGTTATTCTTCTGAGAAAAAAAATGCAGGTGAGGGAGTTAACCA
ATTTCGTAACAAACGGGAAAGTAGAGCGCTATACCGCAGAAGCCCTCCTTGCGCTGCAAGAGGTCAGTTATGAAAAA
TGTCTTATCTCTCTGTTAAGATCCTCTTCATATACATAGTTGCTATTGCTATCGTGAAGTCTTTTTTTTTCTGTTAAT
TGGTCTGGTACTACTTACTAGTCAGGATTTCATATTGCGGTTTTTCCTAGTGGTGTGTAGTTAAAAAGTAGTTTAAT
TGCTTTTAGTTAAAAGGGGTGTTCAGGGCTAAAGATCAACTATGAGAAAACAGAAATTTTCCCAATTCGATACCCGA
CAGCATTATGGCCTGCGCTAATGGAGGTGTTTCCGGGCAAATACTCTAGCCTACCTGGGAAGTACCTTGGGTTGCCC
CTTCATTTCAGGAAAGTAAAAAGGAATGATCTTCAACCTCTAATCGAAAAAATCAACAACAGGCTGGCCTTGCTGGA
AAGGCAAGATGTTGTCCAAGGCTGGTATAGAAACTCTTGTAAAATCGATGCTATCCGCACAACCAATCTACCATCTA
ATGGTTTTTCCACCTCAAAAATGGCTGCTGCAAACAATTGACAAAATACGAAGAAACTTCCTGTGGAGAGGGAGCAA
TCCAGAAGTTTGCAGCGGGGGTCACTGCCTCGTCAACTGGCCCGTAACTTGCCTCCCAAAGAACAAGGGAGGTCTTG
GAATTCTGGACCTTGATCGTTTTGCGAGGGGGCTAAGACTAAGATGGCTGTGGCTACGATGGAAGAGCAAAGATAGG
GCGTGGACTGCCTTGATGCTTCCTTGTGACAAAACTGATGAAGATCTCTTCAATGCTTCCACAACTGTCACGGTAGG
CAATGGAAAGATAGCTGAATTCTGGAATTCTAGTTGGATCCAAGGCCAAGCCCCTAAGAACATTGCGCCAACACTGT

TCAAGAAGGAAAAGAGGAAGAACATCACGGTCGCCAAAGCGCTCACTAACAACAATTGGATTCGTTTATGCTCACCA
TACACGGGTGAGGGGGAGTTTAGAGAGGTCGTCTCTCTTTGGCAGGCCATAGGTAACATGCAAGAGCTTAACGGTTT
GGAAGACAACATCTCTTGGAGATGGACGGCAGATGGGCAGTACAGTGCTAGCAGTGCATACAAAATCCAGTTCGCAT
CCAATTTCACTAAAATGAACCTCTGCCCTATTTGGAAGGCTAAAGTGGAACCGAAATGCCGATTTTTTGCTTGGACA
CTACTTCATAAGAGAATTCTGACTGCCGATAACCTTCATAAAAGAGGTTGCAACTCAGCCTCAGAAACAATTCCCCA
CTTATGCAAGGATTGCCCCTTTAGTAGAGAGGTGTGGAACAAAGTTTTGTCTCGGGCCAACTTTCCTTTACTGACTG
GGTCTCCCAGTGACACTTCTTTGTATGATTGGTGGACGGACATGTGCAGCCTTTGCAGCAGACAGGCAAGAAGAGGT
TTCGACGGTCTGCTATTTCACTTTTGGTGGAACTTATGGCTGGAAAGAAATAACAGAATCTTTCAAAGGCAGCGTAG
AAGTGTAGATCAAGTTGCTCTGGCAGTCAAGGATTATGCTAGTAGCTGAAGTCTAGTTGGTTTGGACTAGTGGTTTT
GTTGCTTTTCTTTTTAATTTCTTTTTAGTTCTTTTTATGTTGTTTTCGTTTCCTTAAGTTGCTTGGAGTCTGTATTA
TCCTCTTTCTTCTAATATAGATCGGAGCGACAAACCTTTTGCCCCTTCCTTTCAAAAAAAAGTTAAAAGGGAATTTA
ACTGCTTTCCTAGTGGTGTAGTTAAAATGGATTTCATATTGCGGCCTTTCCTAGCTTGCTTGCTATTGATTGGACTA
TAGTGATCCAAATGCTGATAACTTTGTCGCTTGTGTAGGCATGGTTAGAGAGCTTAGAGTTTGCATTTATTCAATAC
CTTGAGACTGCATTTCATATACATAAATTATTCATGATTATTTCTTTTCTCTATTTGTTCTGGTTAATTAAGAGTTT
TAGGTTTCCATATTTTTGTACGTGCATCATTTAAATTCTTGTATTGTTTTTCGTTCTTGTCTACAGGCAGCAGAATT
CCACTTGATAGAACTGTTTGAAATGGCGAATCTGTGTGCCATCCATGCCAAGCGTGTCACAATCAGTAAGTTATCAC
TGAGTGAACTCCTTTTTCTCTGTAGCATTACTCCTAATGAATATGTGTGATGCATTTTGGTTGCACGATTCTTTAGT
GATTCTGCTTCAGATGGATATGATAAATCTTGATGTTATTTTGAAGTGGCGAATTGCTTACGAGCGGAAATAGTAAT
GTTCAAATAGCGCAAAGTGCAACTGTTGACTTTTAGTAGGCCATTTATATGGTTTGATTACCAACAAATACGTCAAT
CATATGATTTGATTATCAACAAAGGAATCAGCTATATGGTTTGATTATCAACAAAGGAATCAGCTAGGTTTGCTTAT
CAACATTCAACAAAGGCATCAAGTAATACTCCATCCGTTTCAATTTATAATTCGTTTGACTTTTTTTATCTAAGTTT
GATCGGCTCGACTTATTAAAAAAAATCATAATTATTGTTAATTTTTGTTGTGATATTGTTTAGTATAATATACTTTA
AATGTGACTTTGAGTTTTTCATTTTTTCGCAAAAAAAAATGAATAGGACGAGCCGGTCAAACGTGACACAAAAAAGT
CAAACGAATTATAATTTGGGACACACGGAGTAGTAAATAATGTAACAACTTAGAGAGTGGGACAAAAAAATCTCTAG
TGGTGCTAAATTTAGTTCAGCTTTGTATAAACACAAGCATTGATTGAGAAATCTGACAACTCAAGGATCTGTAGGAA
ATGTGTTACCCTAAATGTTTTCCTTACTGATGCAGTGCAAAAGGACATACAACTTGCAAGGCGTATCGGAGGAAGGC
GTTGGGCATGATATATAATATCCATTCTGATTGCATCATTCTTGTGAATTTGTTTGTAGGAGCTAGACATTAGTGTT
GTTGAATGCTGCATGGTTCCTAATCCTTTTCGCAGTCTAACATCTGTGGAGTTAGTATGTTACATGGCAACAGCTGA
ACATCTGTGGACTATATGGCAACAGCCGAAGATTGTGTCTGTGGGATAACTGGTTGTTTTGGTTGCTCTTCAGTAGT
TTGTTTGCTTCAGGTAACCATGCTGCGAACTATGATGTTTTCATTCTCGGTTTGCTTCAGCTAACCGAGATCGATTC
AGTCTGCAGTATGGACTATGGAGTAAACTGCATGCTGAAACCCGAACCACTGCTGAAACTGCATGCTGAAACCCGAA
CCACTGCTACGGCAGTTGCCAGGATAGCAGGAGGGCCTTTATGCACAGTGGAATTGAGTAGAGAACTGAGTAAACCA
TGGTTCTTTCTCCTTTTGAACTGGAACACACACAGTTGGATCTTGTTTCTCTTCTTAGGCCATTCATCGTGTTTATT
AGGGGTGTAAATGGTACAAATATTATCTGTCCGTATTCGAATTTGATCTATCTAAGAAGGTTGAAATCCGATCCGTA
TTCGAGTCCACCTAGACTTGTCCATCTTCTGGATTGGCCAACTTAATTAATGTATGAAATAAAAGGATGCACACATA
GTGACATGCTAATCACTATAATGTGGGCATCAAAGTTGTGTGTTATGTGTAATTACTAGTTATCTGAATAAAAGAGA
AAGAGATCATCCATATTTCTTATCCTAAATGAATGTCACGTGTCTTTATAATTCTTTGATGAACCAGATGCATTTCA
TTAACCAAATCCATATACATATAAATATTAATCATATATAATTAATATCAATTGGGTTAGCAAAACAAATCTAGTCT
AGGTGTGTTTTGCGAATGCGGC
